# Microporous Immune-Isolating Capsule with Improved Diffusion for Restored Dynamic Bidirectional Hormone Signaling in a Murine Model of Premature Ovarian Insufficiency

**DOI:** 10.64898/2026.04.28.721364

**Authors:** Delaney S. Sinko, Margaret A. Brunette, Despina I. Pavlidis, Monica A. Rionda, Bipasha Ray, Michelle Tong, Suhani Thakur, Brendon Baker, Vasantha Padmanabhan, Ariella Shikanov

**Affiliations:** Biomedical Engineering, University of Michigan, Ann Arbor, MI, USA; Division of Pediatric and Adolescent Gynecology, Eunice Kennedy Shover National Institute of Child Health and Human Development, Bethesda, MD, USA; Department of Chemistry, University of Michigan, Ann Arbor, MI, USA; Department of Materials Science Engineering, University of Michigan, Ann Arbor, MI, USA; Department of Obstetrics and Gynecology, University of Michigan, Ann Arbor, MI, USA; Department of Pediatrics, University of Michigan, Ann Arbor, MI, USA; Department of Molecular and Integrative Physiology, Ann Arbor, MI, USA; Program in Cellular and Molecular Biology, University of Michigan, Ann Arbor, MI, USA

**Keywords:** Thermosensitive Porogens, Immune Isolating Hydrogel, Hormone Restoration

## Abstract

Pediatric cancer survivors treated with gonadotoxic chemotherapy or radiation face lifelong premature ovarian insufficiency (POI), leading to elevated risk of cardiovascular disease, osteoporosis, and metabolic dysfunction. Pharmacological hormone replacement therapy (HRT) cannot replicate the pulsatile, bidirectional signaling of the hypothalamic-pituitary-gonadal (HPG) axis, leaving a critical therapeutic gap. Immune-isolating hydrogel capsules offer a promising strategy for the implantation of donor ovarian tissue without immunosuppression yet they require optimization for human applications. Here, we engineer a microporous immune-isolating capsule by incorporating thermosensitive gelatin microgels as sacrificial porogens. Microfluidic fabrication yielded monodisperse microgels that dissolved at 37°C generating disconnected micropores within a non-degradable poly(ethylene glycol) (PEG) matrix. Critically, the diffusion of FSH-scale analogs (40 kDa) increased by almost two-fold through the microporous capsules relative to nanoporous controls, while antibody-scale molecules (150 kDa) were blocked, demonstrating size-discriminating permeability. In ovariectomized mice implanted with encapsulated ovarian xenografts for 20 weeks, microporous capsules restored dynamic HPG-axis signaling evidenced by elevated levels of estradiol and progesterone, FSH suppression, and fluctuating hormone levels that resembled physiological patterns. Microporosity also improved graft viability, increasing stromal cellularity and reducing follicular apoptosis. These findings support microporous immune-isolating capsules as a platform for physiologically authentic therapy for POI.

## Introduction

Advances in cancer treatment have significantly improved the survival of pediatric cancer patients, with an estimated 85% of children living for 5 years or more after diagnosis^1^. Unfortunately, chemotherapy and radiation have detrimental off-target effects on reproductive organs, causing irreversible depletion of the nonrenewable population of ovarian follicles resulting in premature ovarian insufficiency (POI). Patients with POI have a greater risk of developing chronic conditions, including heart disease, osteoporosis, cognitive dysfunction, growth deficiency, and obesity in adulthood. Consequently, an estimated 50% of adolescent cancer survivors seek follow-up care for hormone imbalances^2^.

The approved treatment to induce puberty in adolescent girls with POI is hormone replacement therapy (HRT), which supplies exogenous estradiol and progesterone to induce breast development, uterine maturation, and menarche^3^.However, exogenously supplied hormones fundamentally cannot replicate the pulsatile, bidirectional signaling of the intact hypothalamic-pituitary-gonadal (HPG) axis; a pulsatile signaling cascade which is driven by follicle-produced estradiol and inhibin regulating the secretion of follicle-stimulating hormone (FSH) and luteinizing hormone (LH) from the pituitary gland. Further, the off-label treatment of young adults with HRT, which was originally designed to treat menopausal symptoms, requires careful monitoring of the timing and dosage of estrogen therapy and leads to suboptimal outcomes and poor patient compliance.

Ovarian tissue-based strategies offer a more physiologically authentic alternative. Ovarian tissue cryopreservation and transplantation (OTCT) can re-establish endocrine function and fertility in patients who cryopreserved tissue prior to cancer treatment. Autografts respond to circulating gonadotropins and secrete ovarian hormones restoring HPG-axis reciprocity before eventual graft exhaustion^4^. Nevertheless, OTCT carries a significant risk of reintroducing malignant cells harbored in the excised tissue, a concern especially acute for patients with hematological malignancies or ovarian metastases. Furthermore, OTCT is only available to patients who banked tissue before gonadotoxic treatment, excluding many children treated before fertility preservation was offered^5,6^.

In contrast to OTCT, transplantation of donor ovarian tissue can restore hormone signaling while addressing both limitations: it avoids re-exposure to the patient’s own malignancies and can benefit patients who never cryopreserved tissue. However, allogeneic tissue is subject to immune-mediated rejection, requiring systemic immunosuppression. To address this, our group developed an immune-isolating encapsulation platform for donor ovarian tissue transplantation (Fig. 1A). The capsule utilizes a dual-layer design: an inner core of proteolytically degradable PEG hydrogel that accommodates tissue growth, surrounded by a nondegradable PEG outer shell whose nanoporous mesh excludes immune cell infiltration and rejection (Fig. 1B). Encapsulated murine ovarian allografts (1 mm^3^) implanted subcutaneously in both naïve and immune-sensitized mice restored endocrine function and resisted rejection for the study duration (4 months)^7^. For translating this approach to human-scale grafts, Brunette et al. scaled up capsule size to accommodate the encapsulation of much larger human ovarian grafts (∼17 mm^3^) to maximize the number of transplanted follicles which are heterogeneously distributed through the ovarian cortex (Fig. 1C)^8,9^. This 17-fold increase in graft volume proportionally increased the diffusion distance that hormones must traverse across the PEG shell. The nanoporous PEG matrix reduces molecular transport by steric obstruction and hydrodynamic drag, effects that scale with path length, larger capsules risk dampening the flux of gonadotropins such as FSH (∼40 kDa) in larger capsules. Insufficient or delayed FSH signaling would impair follicular growth and disrupt the reciprocal estradiol-FSH feedback loop essential for physiological steroidogenesis, undermining the therapeutic benefit of the entire platform. Therefore, introducing controlled microporosity into the PEG shell could selectively accelerate peptide hormone diffusion and benefit folliculogenesis while preserving immune exclusion, provided the pores remain disconnected.

**Figure 1:**
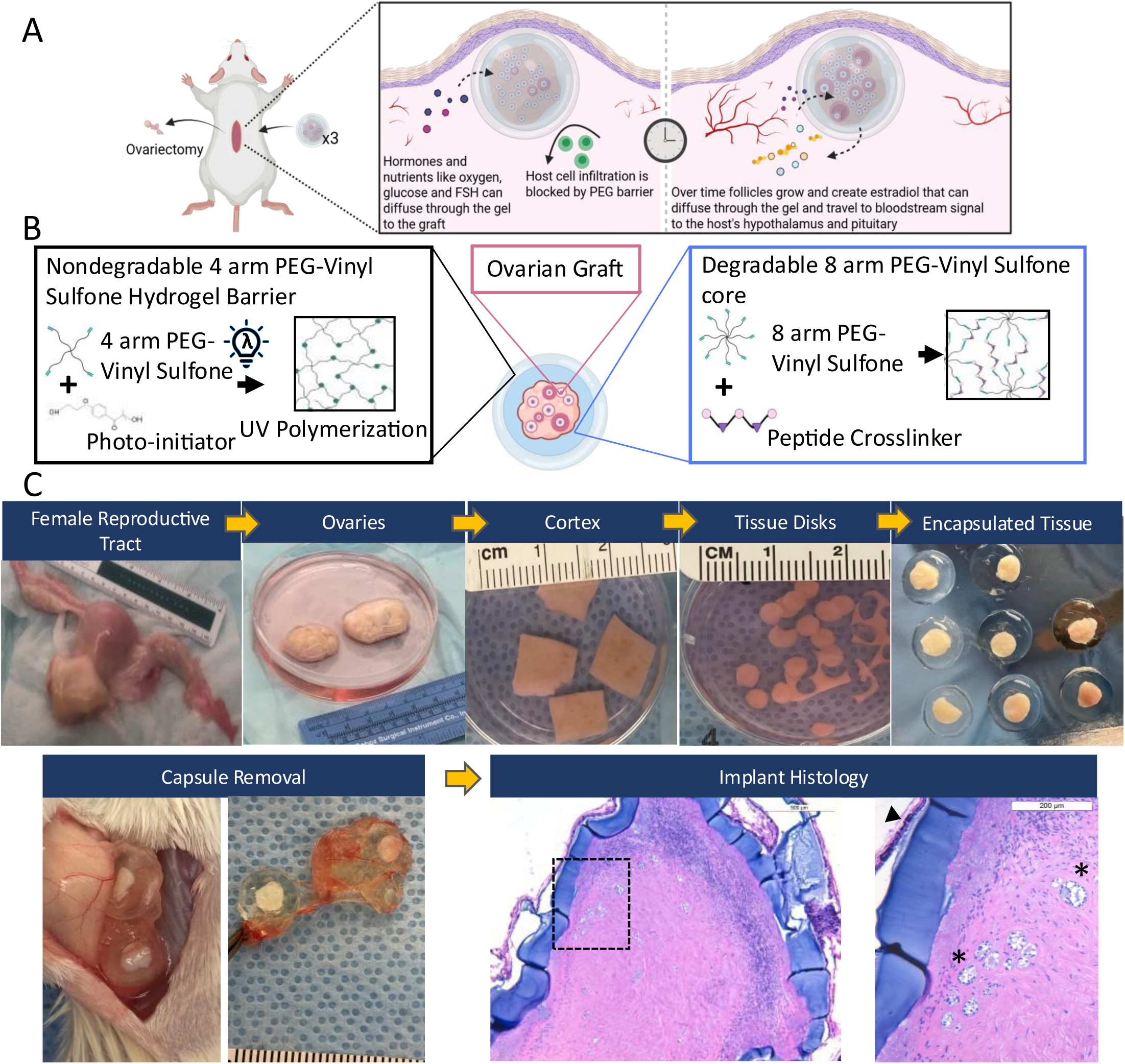
An overview of the immunoisolating capsule composition and human ovarian encapsulation procedure. (A) Schematic overview of implantation and hormone signaling with immune isolating capsule. (B) Hydrogel composition of dual layer immune isolating capsule. (C) Ovarian tissue processing, encapsulation, implantation and removal.

Here, we report the design, fabrication, and in vivo validation of a microporous immune-isolating capsule that enhances the diffusion of peptide hormones, thereby improving the health of encapsulated ovarian grafts. We incorporated thermosensitive gelatin microgels as sacrificial porogens into the nondegradable PEG outer shell; the microgels remained solid during capsule fabrication and tissue encapsulation, then melted at body temperature, leaving behind a network of disconnected micropores. We systematically evaluated capsules containing 1%, 5%, and 10% v/v gelatin (PEG-G+) for the diffusion of peptide hormone-scale and immune molecule-scale tracers, as well as mechanical integrity. We then tested the optimal formulation by transplanting human ovarian xenografts into ovariectomized mice for 20 weeks, monitoring longitudinal HPG-axis hormone profiles and assessing graft histology. This work establishes microporous immune-isolating capsules as a platform for hormonal restoration, providing a physiologically responsive treatment strategy for endocrine restoration in cancer survivors with POI.

## Results and Discussion

### Microfluidic production of gelatin microgels yielded uniform and temperature-sensitive microgels

Using a droplet-generating microfluidic device, we created uniform gelatin microgels in an oil emulsion which retained their shape after transferred to aqueous solution (Fig 2A-C). In the oil phase the diameters measured 26 µm, which then increased to 34.1 µm +/- 2.91 µm after 12 hours of equilibration in aqueous solution and remained unchanged for 120 hours (Fig 2D). Furthermore, the gelatin microgel diameters remained consistent across dozens of production runs, with the majority of diameters between 30 and 36 µm (Fig 2E). To confirm thermo-reversible crosslinking, we heated the gelatin microgels to 37 °C and observed loss of shape followed by complete disappearance of solid microgels in the solution after 10 minutes (Supplemental Figure 1).

**Figure 2:**
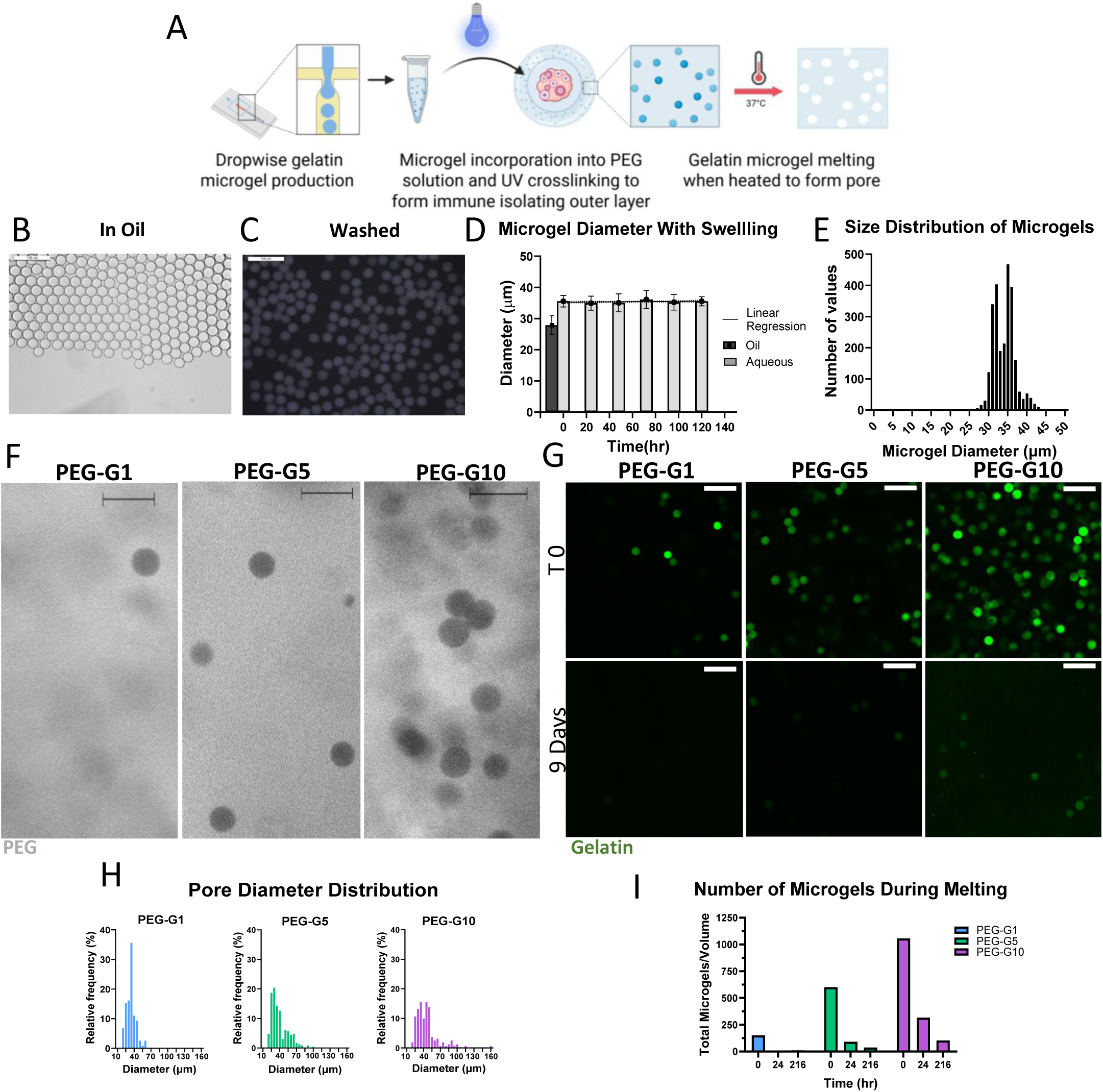
Gelatin microgel porogens had consistent diameters, created pores of corresponding sizes and melted to diffuse out of the PEG gel. (A) Schema c overview of gelatin and PEGG+ composite gel production. Microgels in (B) oil phase and (C) aqueous phase. (D) Microgel diameter after introduc on to aqueous solution up to 5 days; simple linear regression y=0.002531X+35.27, *R*^2^ = 0.001883, F test of slope deviation from 0, nonsignificant p=0.2886.(E) Microgel diameter distribution in aqueous phase over multple product on runs. (F) PEG hydrogel at pore interface. Scale bar represents 50 μm. (G) Fluorescently tagged microgels at crosslinking (T0) and after 9 days meltng at 37°C. Scale bar represents 100 μm. (H) Pore diameter distribution for PEG-G1, PEG-G5 and PEG-G10 gels. (I) Number of microgels counted at crosslinking (t0) and a er 9 days at 37 °C, Chi-square p= <0.0001.

### Gelatin porogens create homogeneously distributed micropores within the nondegradable PEG capsule

To investigate the distribution of the gelatin microgels embedded within PEG hydrogels, we created composite PEG + gelatin gels (PEG-G+) that contained 1% (PEG-G1), 5% (PEG-G5) and 10% (PEG-G10) porogens and quantified fluorescently tagged gelatin microgels throughout the x, y and z planes using confocal microscopy (Fig 2F). As expected, single gelatin microgels created pores with diameters of 35-40 µm, with larger pores observed due to aggregation, the frequency of which positively correlated with the volume fraction of microgels incorporated. The percentage of pores ranging between 20-40 μm was 84.7%, 70.9% and 51.3% for PEG-G1, PEG-G5 and PEG-G10 gels, respectively (Fig 2H). The lowest percentage of larger pores measuring 40 – 50 μm that formed due to microgel aggregation in PEG-G1 gels (15.3%), compared to 19.1% in PEG-G5 and 35.6% in PEG-G10. Larger aggregates of microgels measuring between 60 and 120µm in diameter were observed at 10.0% and 12.5% of pores in the PEG-G5 and PEG-G10 gels, respectively. Moreover, in the PEG-10G gels some pores reached 160 μm, which was not present in any other condition. This positive correlation between the volume fraction of gelatin microgels and the percentage of larger pores suggests that porogen content above 10% in the PEG-G+ gels could likely create large, connected pores, increasing the risk of host cell infiltration and potentially compromising the immune-protective properties of the PEG shell.

We then investigated the kinetics of gelatin melting and diffusion by incubating composite PEG-G+ gels containing fluorescently tagged gelatin microgels at 37°C to mimic body temperature (Fig 2G). The initial number of clearly defined spherical microgels counted within a region of interest (ROI) of the gel was 150 (PEG-G1), 601 (PEG-G5) and 1056 (PEG-10) (Fig 2G). After 24 hours at 37°C the number of microgels decreased across all conditions to 4 (PEG-G1), 91 (PEG-G5) and 316 (PEG-G10) remaining microgels. Finally, after 9 days at 37°C, only PEG-G10 gels retained 10% of the initial load of measurable microgels, while microgels in PEG-G1 and PEG-G5 were undetectable. A chi-squared analysis revealed a statistically significant relationship between melting time and microgel number (p < 0.0001). The melting occurred exponentially, representing a first order kinetic release profile of gelatin during diffusion out of the matrix. This suggests the rate of gelatin removal was not dependent on heating, but the concentration of gelatin in the system. Due to this the melting and dissolution of gelatin out of the PEG matrix is the slowest in the PEG-G10 gels.

### Micropores significantly enhanced diffusion of peptide hormone-scale molecules while preserving exclusion of immune molecules

Next, we compared the diffusion of molecules ranging from 4 to 150 kDa through PEG-G+ gels with increasing micropore concentrations to determine how biological molecules of different sizes are affected by microporous gels. We used a broad range of molecular masses of dextran to represent biological molecules; small molecules such as glucose were modeled with 4 kDa, peptide hormones were represented by 40 kDa molecules, and large proteins, such as antibodies, were modeled with 150 kDa. The mass transfer, or the percentage of molecules that diffuse through the gel, of 4 kDa dextran in the PEG-G+ gels was not affected by the increasing micropore density and did not differ from the nanoporous controls, reaching 60-75% after 96 hours across all condtions (Fig. 3A&3D). The addition of micropores, however, significantly increased the diffusion of larger molecules; the diffusion of 40 kDa dextran almost doubled, increasing from 17.2% in the unmodified controls to 29.8% in the PEG-G10 gels (p=0.0001), 25.5% in the PEG-G5 gels (p=0.0002) and 24% in the PEG-G1 gels (p=0.0006) (Fig 3B & 3E). Finally, as expected, the diffusion of 150 kDa dextran was the slowest of all the molecules and reached 11.2% in control gels but increased significantly to 17.8% in only PEG-G10 gels (p=0.0084) (Fig 3C & 3F). In summary, the addition of micropores increased the diffusion of peptide hormone scale molecules (40kDa) in PEG-G+ gels by 173%, with no significant changes in the diffusion of small molecules (less than 4 kDa). Across conditions, PEG-G1 and PEG-G5 gels demonstrated a significant increase in diffusion for peptide hormone scale molecules but remained unchanged for large molecules compared to nanoporous controls. In contrast, the increased diffusion of large immune molecules in PEG-G10 gels may have resulted from microgel aggregation, suggesting the potential for large interpenetrating pores that could compromise immune isolation when implanted in vivo. These findings indicate that the main risk for this technology is the potential aggregation of micropores, especially at a higher micropore content, which may create interconnected pathways that permit cellular infiltration and undermine immunoisolation. Importantly, PEG-G5 gels achieved similar improvements in diffusion but with fewer large pores, therefore balancing transport and barrier integrity.

**Figure 3:**
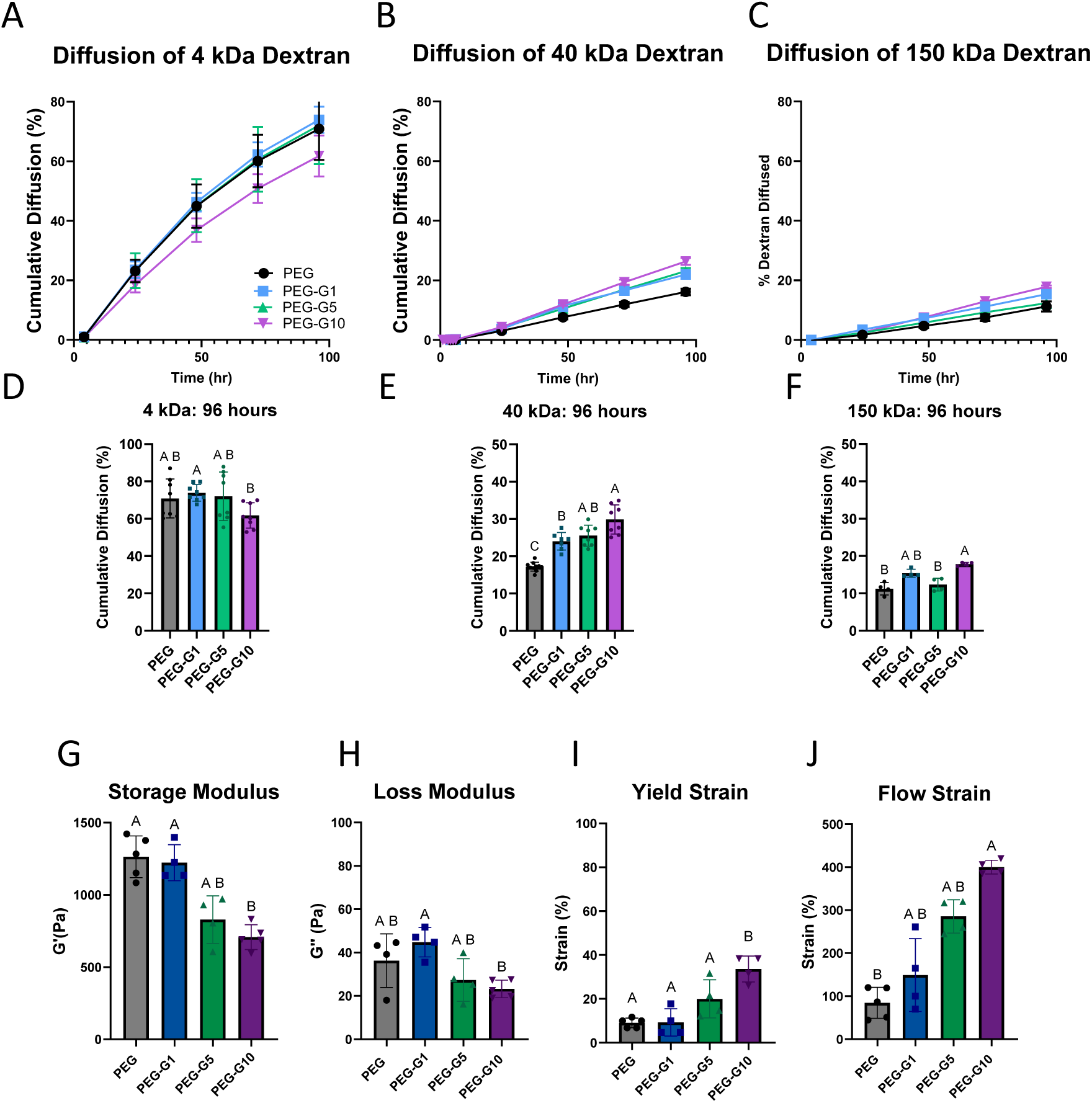
Diffusion of small protein scale molecules increased for PEG-G+ gels while mechanical profiles demonstrated decreased elasticity but increased distensibility. Diffusion of (A) 4 kDa dextran, (B) 40 kDa Dextran, and (C) 150 kDa dextran across PEG and PEG-G+ gels over me. Cumula ve diffusion of (D) 4 kDa dextran, n=8, ANOVA with Brown- Forsythe + Welch tests letters dictate statistical difference (p<0.05), (E) 40 kDa dextran n=8, ANOVA with Brown- Forsythe + Welch tests letters dictate statistical difference (p<0.05) and (F) 150 kDa dextran after 96 hours, n=4 ANOVA with Kruskal-Wallis post hoc test, letters dictate significance (p<0.05). (G) Storage modulus at 37°C and (H) loss modulus at 37°C. (I) Yield Strain and (J) flow strain at 37°C n=4-5 ANOVA with Kruskal Wallis post hoc test, letters dictate statistical difference (p<0.05).

Using Fick’s Law, we calculated the diffusivity of various biological molecules from cumulative diffusion data, time, and path length to estimate the diffusion time of a single molecule across the immune-isolating capsule^10^. To travel the thickness of the capsule (2mm), small molecules (4 kDa) would take less than an hour regardless of whether the micropores were present or not. Molecules in the peptide hormone range (40 kDa) would take 12.5 hours to pass through the control nanoporous gels, compared to 4.7 hours to travel through the PEG-G10 gel capsule, representing a 150% increase (Supplemental Figure 3). Therefore, micropores within a nanoporous gel accelerate the flux of biological molecules across the hydrogel and improve the dynamic timing of bidirectional hormone communication.

### Micropore inclusion reduced gel elasticity while enhancing the strain required to cause polymer network breakdown and solid —to— fluid transition, preserving mechanical integrity

To investigate the effect of micropores on the mechanical profile of the composite PEG-G+ hydrogels, we performed an increasing oscillatory strain test on a parallel plate rheometer. Storage and loss moduli were calculated over the linear viscoelastic region of the strain sweep to measure the elasticity and viscosity of the gels at 37°C^11^. The storage modulus decreased with the addition of micropores from 1264 Pa for nanoporous gels to 708 Pa in the PEG-G10 gels (p=0.0124) (Fig. 3G). Meanwhile, the loss modulus of microporous gels remained similar compared to control nanoporous PEG gels (Fig. 3H). This data suggests that PEG-G+ gels are less elastic under moderate levels of strain which is expected as the micropores disrupt the covalently bound polymer chain network.

To understand whether micropores negatively affect the structural integrity of the PEG barrier we compared the strain at shear thinning (yield strain) and solid-to-fluid transition (flow strain) of PEG-G+ gels to nanoporous gels^12^. In PEG-G+ gels the linear viscoelastic region of microporous gels was extended, as demonstrated by an increased yield strain. The average strain required to yield the PEG-G10 gels was 33.5% compared to 9.06% for nanoporous PEG gels (p=0.0041) (Fig. 3I). Generally, yield strain indicates an initiation of plastic deformation or shear thinning behavior that is defined by the force required to cause alignment of polymer chains, and the breaking of crosslinking bonds. Therefore, an increased yield strain indicates the microporous gel is able to withstand larger forces before nonrecoverable damage is initiated on the system. Furthermore, microporous PEG-G+ gels exhibited an extended region of viscoplastic deformation with a flow strain of 400% (PEG-G10), 285% (PEG-G5) and 149% (PEG-G1) compared to 84.2% in nanoporous controls (Fig. 3J). The flow strain marks the end of plastic deformation, and complete structural breakdown where the material is no longer solid. Therefore, the yield data suggests that the inclusion of micropores increases the distensibility of the gels, but does not prompt mechanical failure at lower stresses, implying that the gels are still able to withstand the forces required for implantation.

### Human ovarian tissue encapsulated in PEG-G+ capsules supported folliculogenesis, improved stromal cell survival and restored hypothalamus- pituitary signaling in ovariectomized mice

To investigate the effect of microporosity of immunoisolating capsules on graft health, we encapsulated human ovarian tissue in PEG-G5 and PEG-G10 modified capsules and implanted them subcutaneously in ovariectomized immune-deficient mice for 2, 10 or 20 weeks. Nanoporous PEG capsules served as unmodified controls. Histological analysis confirmed the presence of follicles within the grafted tissue at all time points, and increasing follicle maturity with time in vivo suggesting active folliculogenesis in the tissue. Most of the follicles present in the grafts at the time of transplantation are primordial and primary follicles, as they constitute the follicular pool in the cryopreserved cortex. The progression from the primordial and primary stages to the secondary and antral stages takes months in human follicles. Thus, in the grafts retrieved after two weeks post-transplantation, we identified mostly primordial and primary follicles (Fig. 4A). After long term implantation (10 weeks), growing follicles with multiple layers of granulosa cells surrounding the oocyte and larger follicles with fluid-filled antrum were observed in the grafts (Fig. 4B). Concurrently, significant populations of primordial and primary follicles were still present in tissue, suggesting stepwise activation and growth leading to an extended hormone production and endocrine function.

**Figure 4:**
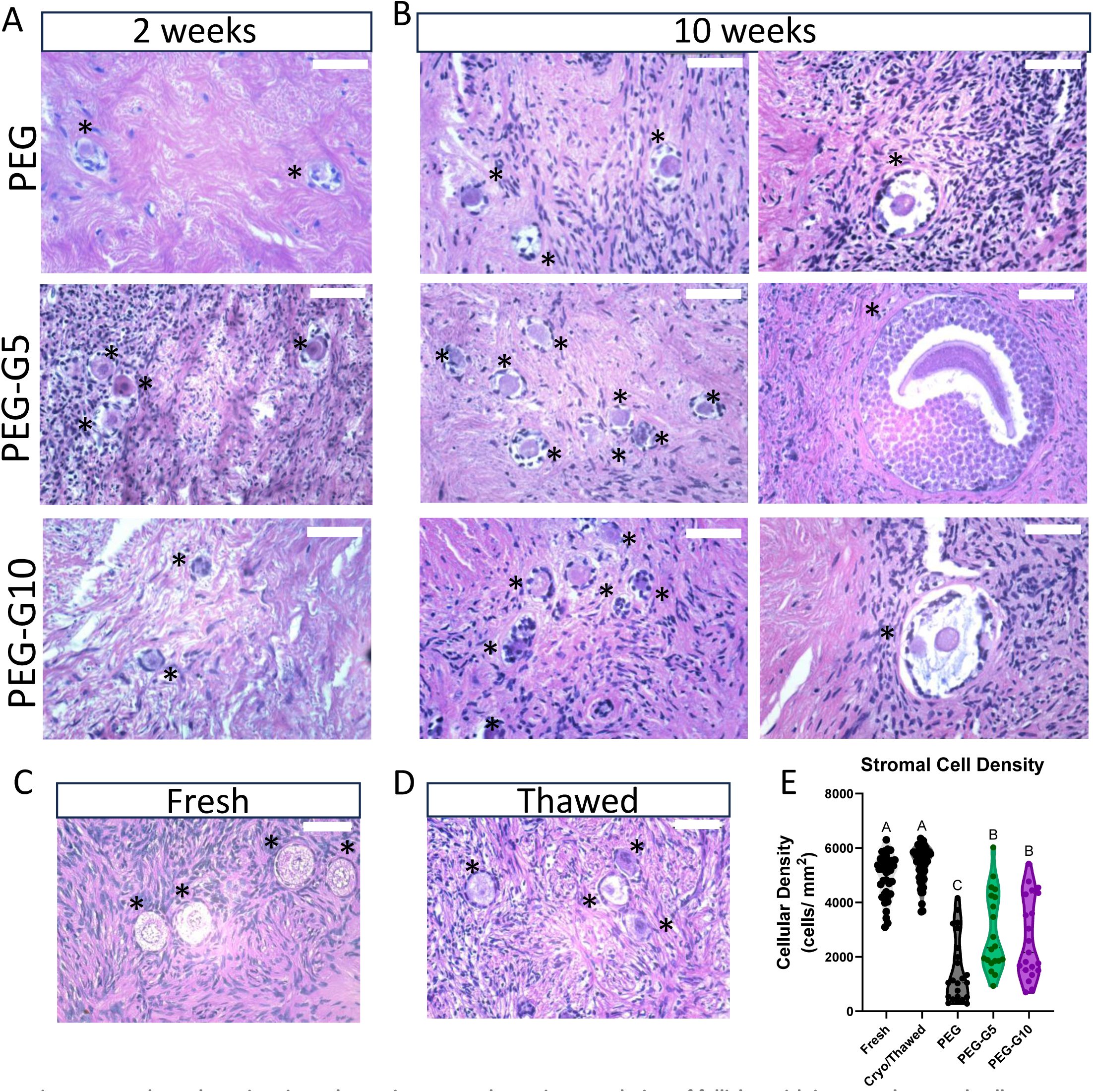
Implanted ovarian tissue has quiescent and growing population of follicles with increased stromal cell populations. Representative histology images of implants after (A) 2 weeks, and (B)10 weeks. Scale bar represents 50 μm, asterisks indicate follicles. Representative (C) fresh and (D) thawed ovarian tissue. (E)Stromal cell density of implant at 20 weeks post implantation, lognormal ANOVA with Brown-Forsythe and welch post hoc test, letters dictate statistical difference (p<0.05).

Next, we quantified the stromal cells by measuring the cellular density around the follicles in the implants as an indicator for tissue health. Human ovarian tissue is heterogeneous, therefore, we expected to find significant variation in stromal cell densities. Cellularity of fresh tissue from the three donors ranged between 3,090 to 6,299 cells per mm^2^, with an average of 4,919 cells per mm^2^(Fig 4C). The cellularity in tissues that were cryopreserved, thawed and immediately fixed had the same cellularity as fresh tissue (Fig 4D&4E). After long-term implantation, the number of stromal cells decreased the most in the unmodified controls, reaching 1,498 cells per mm^2^ (p<0.001), compared to 2,906 (PEG-G5, p<0.001) and 2,695 (PEG-G10, p<0.001) cells per mm^2^ in PEGG+ gels (Fig 4E). This data suggests that all encapsulated tissue experienced a decrease in stromal cell density, likely due to the avascular environment, but in the presence of micropores, more stromal cells survived.

The restoration of endocrine function in ovariectomized mice was confirmed by the estrous cyclicity and systemic levels of hormones, FSH, estradiol and progesterone. In general, ovariectomy leads to a cessation of estrous activity, a drastic decrease in systemic levels of estradiol and progesterone, and a significant rise in the levels of FSH in the absence of negative feedback from ovarian hormones. As expected, after ovariectomy, the estrous activity ceased, confirmed by persistent diestrus in all mice. Estrus activity was restored in 70% of mice following implantation of encapsulated ovarian grafts across all PEG-G+ and PEG conditions, along with expected physiological changes associated with hormone fluctuations, such as vaginal opening (Supplemental Figure 6). After the implantation of encapsulated ovarian grafts in ovariectomized mice estradiol levels increased from undetectable, or below 5 pg/mL after ovariectomy to an average of 200 pg/mL after 10 weeks and to 300 pg/mL after 20 weeks (Fig. 5A). Corresponding serum FSH decreased to physiologic levels measured in intact mice, ranging between 14-18 ng/mL for all the conditions compared to 80ng/mL in ovariectomized control mice (Fig 5B). Furthermore, after 10- and 20-weeks progesterone levels reached an average of 12-15 ng/mL across all groups (Fig. 5C). The increased levels of circulating estradiol and progesterone paired with decreased levels of FSH confirmed that the grafted human ovarian tissue fully restored the physiological feedback between the hypothalamus, pituitary, and ovarian tissue across all conditions. The absence of statistically significant differences in average hormone levels between microporous and nanoporous capsules most likely reflects the inherent biological heterogeneity of human ovarian tissue rather than a lack of biological effect. Follicle density is variable across donors and even within regions of the same ovary, and the lack of non-destructive methods for quantifying follicle content prior to implantation introduces substantial variance, reducing statistical power.

**Figure 5:**
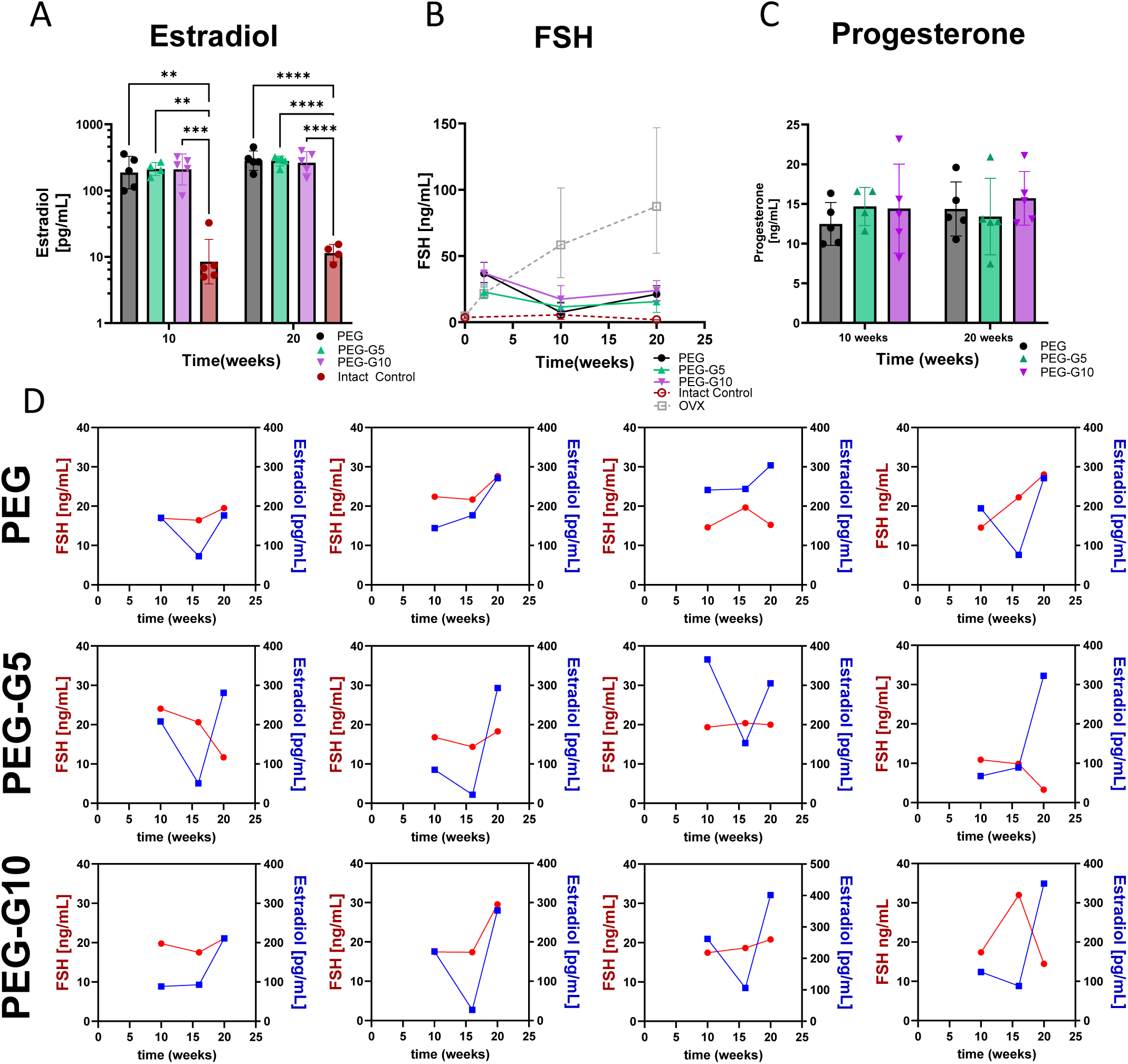
Terminal serum levels demonstrated restored ovarian hormones and HPG axis regulation, and dynamic hormone fluctuations over time. (A) Pooled estradiol levels 10 and 20 weeks post implantation, n=4- 5, 2way ANOVA stars dictate statistical significance. (B) Pooled FSH levels at 10 and 20 weeks, n=4-5. (C) Pooled serum progesterone levels at 10 and 20 weeks post implantation. (D) Estradiol and FSH levels for individual mice at 10, 16 and 20 weeks for PEG and PEG-G+ gels n=4-5. OVX denotes ovariectomized.

**Figure 6:**
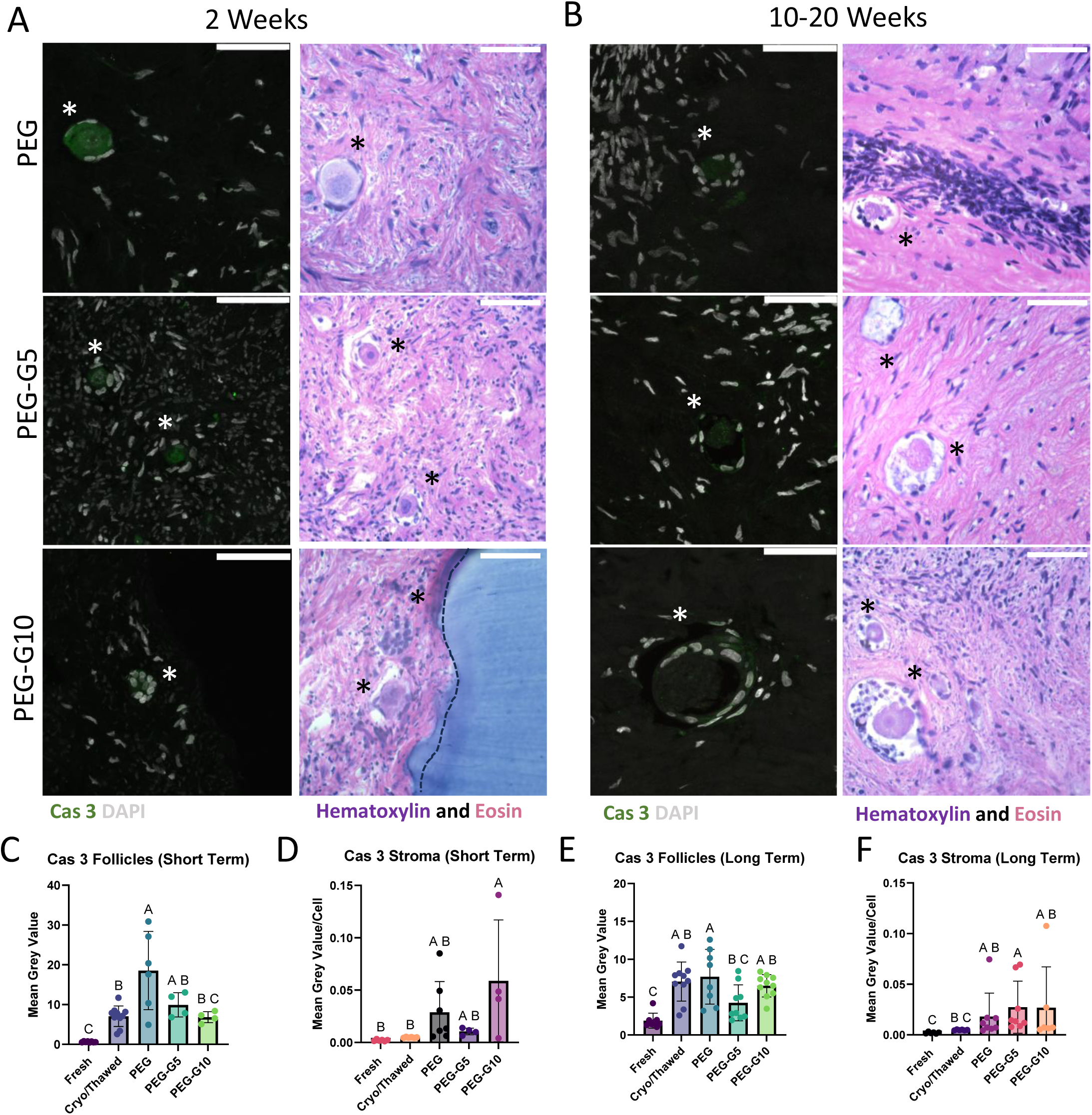
Microporous encapsulation decreased follicular apoptosis initially after implantation. Cleaved caspase staining and corresponding histology images of implants after (A) 2 weeks of implantation and (B) 10-20 weeks post implantation. Scale bar represents 50 μm. Mean grey value of cleaved caspase staining in (C) follicle n = 4-10, ordinary one-way ANOVA with Tukeys post hoc test (D) surrounding stroma at 2 weeks n=4-7, ordinary one-way ANOVA with Tukeys post hoc test. Mean grey value of cleaved caspase 3 in (E) follicle, n= 8-10, ordinary one way ANOVA with Tukeys post hoc test and (F) surrounding stroma at 10 and 20 weeks (pooled), n = 6-10 nonparametric ANOVA with Kruskal-Wallis test. Scale bar represents 50 μm. Letters dictate statistical differences (p<0.05)

To determine whether micropores enhanced bidirectional hormone exchange between the graft and host, we monitored longitudinal hormone profiles in individual animals at 10, 16, and 20 weeks. Strikingly, mice demonstrated fluctuating levels of estradiol and FSH throughout this extended period, recapitulating the physiological dynamic reciprocity of the HPG axis. Estradiol levels oscillated between values greater than 300 pg/mL and as low as 100 pg/mL (Fig 5D), while FSH fluctuated correspondingly within the range of 10–30 ng/mL across all encapsulated conditions (Fig. 5D). Critically, the most pronounced hormonal fluctuations were observed in mice receiving PEG-G10 capsules compared to nanoporous controls, directly implicating micropore - mediated diffusion in enhancing the timing and amplitude of bidirectional hormone signaling. These findings demonstrate that the addition of micropores does not merely increase passive hormone transport, but enables a more physiologically responsive, dynamic exchange between the encapsulated graft and the host HPG axis, a capacity that exogenous HRT fundamentally cannot replicate.

Finally, we assessed whether addition of the micropores compromised gels’ ability to withstand host cell infiltration for 20 weeks in vivo. We analyzed the retrieved grafts for the presence of red blood cells, which would indicate cellular infiltration from the host and compromised immune isolation. In control PEG capsules, only 1capsule had cellular invasion, consistent with our prior results^13^. As expected, PEG-G10 gel condition had the greatest number of gels with host cells invading the grafts, because a greater number of pores increased the risk of interconnected larger channels. To further assess graft health, we analyzed only intact, fully isolated ovarian tissue, as the presence of blood vessels would have influenced the results.

### Follicle and stromal cell health improved in tissue encapsulated in micropore-containing hydrogel capsules

Next, we quantified cell death marker, cleaved caspase 3 (Cas3) to determine whether increased diffusion decreased apoptosis in the implanted tissue^14^. Fresh ovarian tissue had low levels of Cas 3 with an average intensity of 0.6113, which increased to 7.062 in cryopreserved samples. After 2 weeks, implanted follicles in nanoporous control hydrogels demonstrated a statistically significant increase in Cas3 proteins with a mean grey value of 18.54 (p=0.0021) (Fig. 5A&5C) compared to cryopreserved controls, whereas follicles encapsulated in PEG-G+ gels increased only to 9.927 (PEG-G5) and 6.8245 (PEG-G10) and were not significantly higher than cryopreserved controls. After long-term implantation (10 and 20 weeks), the levels of Cas3 decreased across all groups and closely resembled cryopreserved controls. The levels of Cas3 in follicles in the PEG controls and PEG-G10 remained elevated compared to fresh controls with average intensities of 7.698 (p=0.0013) and 6.501 (p=0.0031), respectively (5B,5E&5F). Similar trends were observed in the quantification of mixed lineage kinase domain like protein (MLKL) which executes damage-initiated necroptosis (Supplemental Fig. 7). This data suggested that most events associated with cell distress occur at the earlier time points, and follicles that survive the initial shock exhibit low levels of cell death markers and stress. Elevated levels of Cas3 in the long-term conditions are likely due to the avascular state of the tissue; nevertheless, we observe follicular growth and hormone production in the encapsulated tissue.

Furthermore, to assess stromal cell apoptosis levels Cas3 was quantified in the stroma directly surrounding the follicles. In fresh tissue the stroma exhibited low levels of Cas3 with that doubled in cryopreserved/thawed tissue. After 2 weeks in vivo, the levels of Cas 3 in the stroma around the follicles increased slightly compared to fresh and cryopreserved controls. The tissue in PEG-G10 gels exhibited the highest expression at 0.0588 (p=0.0229) mean grey value per cell compared to fresh controls (Fig. 5D). Interestingly, PEG-G5 gels had the lowest Cas 3 levels. At long-term timepoints, PEG-G10 gels had slightly increased values with an intensity of 0.025 (p=0.0129) mean grey value per cell compared to fresh controls, but all groups were not significantly different from cryopreserved ovarian tissue (Fig. 5F). The relative stability of the stromal levels of Cas 3 over many weeks of encapsulation suggests the tissue did not experience elevated damage as a result of encapsulation.

After encapsulation in microporous immunoisolating capsules, PEGG+ gels primarily improved outcomes at the cell and tissue scales by increasing stromal cell survival and reducing follicular apoptosis. These improvements in tissue health were likely due to the flux of FSH to the graft. FSH is a peptide hormone essential for folliculogenesis and steroidogenesis, as it stimulates proliferation and maturation of granulosa cells by activating the AKR/NFκB pathway, which blocks caspase-mediated apoptosis ^15–17^. With a molecular mass of 35 kDa, FSH is susceptible to slowing within the nanoporous immunoisolating capsule, which would disrupt the kinetics of hormone signaling. The microporous architecture accelerated FSH transport across the capsule wall, likely delivering a pro-survival rescue signal to the graft during the critical early post-implantation period. Consistent with this, follicles in control nanoporous gels had increased levels of Cas3 after 2 weeks compared to follicles encapsulated in microporous gels. Since stromal cells support follicle growth and maturation through paracrine signaling and extracellular matrix deposition, their survival represents a compounding benefit: a healthier stromal environment sustains folliculogenesis over the long term, amplifying the downstream hormone production that the aggregate ELISA measurements only partially capture^18^.

### Conclusions

Cell-based hormone restoration, in contrast to HRT, inherently adapts to the biological variation between individuals, offering a self-regulating approach to hormone restoration that static replacement therapies cannot replicate. In this study, we introduced micropores into a nanoporous PEG hydrogel-based immunoisolating capsule and evaluated whether increased diffusion could improve the survival and function of human ovarian grafts implanted in ovariectomized mice. Here, we adapted this approach to establish a microporous platform that enhances diffusion without compromising the immune-isolating properties. Micropore-enabled enhancement of bidirectional diffusion accelerated hormone transport, increasing FSH signaling to the graft, in turn, translating into measurable biological benefits, including reduced follicular apoptosis and increased stromal cell survival. Further, microporous encapsulation likely accelerated both the inward transport of gonadotropins to the graft and the outward communication of ovarian steroids to the pituitary, therefore tightening the feedback loop that underlies normal endocrine physiology.

While this study yielded exciting results, there are several limitations that inform future work. First, the murine model, while well-validated for HPG-axis signaling and a practical platform for proof-of-concept encapsulation studies, has a lifespan far shorter than the human reproductive period, constraining our ability to assess graft longevity and the sustainability of endocrine function over clinically relevant timescales. Second, the xenogeneic nature of this model presents a challenge when assessing the effect of the immune system on this technology. Addressing these limitations will require studies in humanized immunocompetent models, which would permit investigation of donor-recipient immune interactions under more physiologically relevant conditions, and ultimately in large animal models that better approximate human reproductive timescales and anatomy. Together, these future studies will define the immune risk profile of microporous capsules and establish the boundary conditions for long-term graft health.

Lastly, beyond their role as porogens, the incorporation of sacrificial microgels present a platform for delivery for further engineering of the implant microenvironment. As the microgels dissolve, they create a transient and characterizable window for delivery of bioactive molecules, such as immune- modulating cytokines, angiogenic factors or follicle activating signals. This is particularly advantageous at the early implantation period, when immune surveillance is heightened. Exploiting this delivery window to further control how the implant interfaces with the body would address principal bottlenecks that currently limit the performance of the immune isolating capsule, moving this platform closer to a clinically viable endocrine restoration therapy for cancer survivors with POI.

## Methods

### Microfluidic Fabrication of Gelatin Microgels

A droplet-forming two-inlet junction microfluidic device was used to create uniform gelatin microgels based on a previously established chip system^22^. First, a silicon wafer negative was fabricated using lithography methods then a PDMS mold of the silicon negative was created and cured to produce microfluidic devices. Biopsy punches (0.5mm, 0.74mm and 1.2mm) were used to cut the needle entry holes and outlet port. Finally, the PDMS devices and bottom coverslip were plasma etched for 1 minute and bonded. The devices were incubated at 60 ⁰C overnight to confirm bonding.

Porcine skin gelatin (Sigma-Aldrich, G2500) was dissolved in 37⁰C Dulbecco’s Phosphate Buffered Saline (DPBS) (Fisher Scientific, 14-040-133) at a concentration of 2.5% w/v and was run through the aqueous inlet of the device. A solution of 1% fluorosurfactant (RAN Technologies, 008-FluoroSurfactant-5wtH) in Novec 7500 oil (Best Technology Inc, 98-021-2928-5) was run through the oil inlet. Flow rates were modulated until a stable flow of approximately 30 µm droplets was being pinched off at the junction. The resulting emulsion was transitioned into the aqueous state and stored in DPBS at 4° C.

### Formation of PEG-G+ Hydrogels

A non-degradable poly(ethylene glycol) (PEG) hydrogel with embedded gelatin microgels was adapted from a previously defined protocol^23^. In short, 4-arm PEG-VS (Jenkem, PEG-Vinyl Sulfone hexaglycerol) was dissolved in DPBS with 0.4% Irgacure 2959 (Millipore Sigma, 410896) and 0.1% N-vinyl-2-pyrrolidone (NVP) (Millipore Sigma, V3409), creating a final concentration of 5% w/v PEG. The concentration of gelatin microgels in DPBS was calculated using a hemocytometer and the appropriate volume of solution was dispensed to ensure the correct volume of gelatin microgels. After centrifugation the gelatin microgels formed a pellet, and the supernatant was aspirated, leaving only the microgels behind. PEG solution was added to the gelatin and mixed thoroughly by pipetting up and down. The resulting gel precursor was added to a PDMS mold and crosslinked with UV light (Lumen Diagnostics, 200W Lamp S2000) 30mV/cm^2^ for 45 seconds.

The PEG-G+ gels were characterized using image analysis. The PEG and gelatin gels were labeled with fluorescent probes conjugated to CF640R maleimide and CF488A succinyl ester (Biotium, 50-196-4889), respectively. Various timepoints of PEG-G+ gels were imaged using confocal microscopy and the Weka image classifier of Image J was used to analyze the porosity, number of pores and distribution in the Z stacks of gels.

### In Vitro Diffusion Characterization

PEG-G+ gels were crosslinked in a transwell (Corning, 07-200-154) and swelled for 4 days in DPBS at 37°C. Then 150 μL of FITC-Dextran solution (Milipore Sigma, FD2000S)(2 µM) was added on top of the gel and the transwell was placed in a well containing DPBS. An empty transwell containing 150 μL of FITC-Dextran solution served as a positive control. The transwell was moved hourly to a new well for the first 5 hours and daily up to 5 days where 100 μL of the DBPS solution was sampled. After 5 days the concentration of the 488 FITC was measured in all samples using a plate reader (Biotek, Synergy, H1 Multimode Microplate Reader). The amount of the dextran that diffused through the gel was interpolated using standard curves with known concentrations. A modified Fick’s law was used to calculate the effective diffusion coefficient for each sample using the diffusion data and test parameters^10^. To estimate the time required for biological molecules to diffuse through the PEG-G+ gels, the hydrodynamic radii of the molecules were used with the gel-estimated diffusivity.

### Rheology of PEG-G+ gels

PEG-G+ gels (2 mm thick) were crosslinked swelled at 37°C for 5-7 days to allow for melting and diffusion of the gelatin. The gels were biopsy-punched into 8mm disks and loaded into a Discovery Hybrid parallel plate rheometer-2 (TA Instruments, DHR-2) while maintained at 37°C. Frequency and time sweeps were performed to define parameters within the linear viscoelastic region. Then, a 0.1-1000% strain sweep was performed at a frequency of 1 Hz. Yield strain was defined as the strain at which a 5% decrease from the storage modulus of the linear viscoelastic region was observed, and flow strain was determined using the crossing over point of the G’ and G’’ moduli.

### Ethical Approval Process and Tissue Procurement

Whole human ovaries were obtained from deceased donors for non-clinical research through the International Institute for the Advancement of Medicine (IIAM) and associated Organ Procurement Organizations (OPOs). IIAM and the associated OPOs comply with state Uniform Anatomical Gift Acts (UAGA) and are certified and regulated by the Centers for Medicare and Medicaid Services (CMS). These OPOs are members of the Organ Procurement and Transplantation Network (OPTN) and the United Network for Organ Sharing (UNOS) and operate under standards established by the Association of Organ Procurement Organizations (AOPO) and UNOS. Informed, written consent was obtained from the deceased donors’ families prior to tissue procurement for the tissue used in this study. A biomaterial transfer agreement is in place between IIAM and the University of Michigan that restricts the use of the tissue to pre-clinical research and does not involve the fertilization of gametes. The use of deceased donor ovarian tissue in this research is categorized as “not regulated” per 45 CFR 46.102 and the “Common Rule,” as it does not involve human subjects and therefore complies with the University of Michigan Institutional Review Board (IRB) requirements.

Ovarian tissue samples were procured from 3 healthy deceased donors of reproductive age (33, 23 and 22 years old) with follicle densities ranging from 84.04-119.30 follicles/mm^3^ (Supplemental Fig. 5). The tissue was processed into 10mm x 10 mm x 1mm squares, which were then slow frozen^9^. Briefly, squares were equilibrated in 900 μl of freeze media consisting of 10% Serum Protein Substitute (Origio, ART-3010), 0.75M dimethyl sulfoxide (DMSO) (SigmaAldrich, D2653), 0.75M ethylene glycol (SigmaAldrich, 102466), 0.1 M sucrose (SigmaAldrich, S1888) in Quin’s Advantage media (Origio, ART-1024) at 4°C for 30 minutes before being slow cooled using Cryologic Freeze Control System (Cryologic, Australia). Samples were cooled from 4 °C to -9 °C at a rate of -2 °C /min, equilibrated at -9 °C, seeded to induce ice crystal formation, held for 5 minutes at -9 °C and cooled to -40 °C at a rate of -0.3 °C / min. Samples were stored at -196 °C.

### Ethical Use of Animals Statement

All procedures performed using animals were in accordance with standards dictated by the Institutional Animal Care & Use Committee and the University of Michigan protocol PRO00011284.

### Formation and Implantation of PEG-G+ Immune Isolating Capsules

Slow frozen ovarian cortex pieces were thawed in a 37°C water bath and removed just after ice had melted. The tissue was then maintained at room temperature and washed of cryoprotectants using a 10 minute washes in the following media concentrations: (1) 0.5 M Ethylene Glycol, 0.5 M DMSO, 0.1 M Sucrose, 10% Serum protein substitute in Quins Advantage Media (2) 0.25 M Ethylene Glycol, 0.25 M DMSO, 0.1 M Sucrose, 10% Serum protein substitute in Quins Advantage Media, (3) 0.1 M Sucrose, 10% Serum protein substitute in Quins Advantage Media and (4) 0.1 M sucrose in 10% Serum Protein Substitute. The tissue was then biopsy punched into 4 mm disks and encapsulated in 40 μL of degradable hydrogel layer crosslinked using 8-arm PEG-VS (Jenkem Technology, PEG-Vinyl Sulfone hexaglycerol) with a plasmin sensitive peptide sequence YKNS (Ac-GCYK↓NSGCYK↓NSCG, Genscript) at a 1:1 ratio of -Vs to thiol groups. Then the 8-arm PEG hydrogel encapsulated tissue was placed in a 120 μL droplet of UV cross linkable PEG-G+ hydrogel detailed above and crosslinked under UV for 45 seconds flipping at a midpoint to allow even crosslinking. Implants were stored in Leibovit’s L-15 medium (Thermofisher, 11415064) at room temperature until implantation.

Female immunocompromised mice (NOD.Cg-*Prkdc^scid^ Il2rg^tm1Wyl^*/SzJ, Jackson Laboratories) aged 6 to 8 weeks were ovariectomized and subcutaneously grafted with 3 capsules. Briefly, a dorsal central incision was created, followed by two lateral incisions in the muscle. The ovaries were cauterized to extract, followed by the closing the peritoneal muscle, insertion of the implants in the subcutaneous space, and final closure. Two mice per condition were grafted for 2 weeks and five mice per condition were grafted for 10 and 20 weeks. To monitor estrous cyclicity vaginal cytology was performed daily and blood was collected every 2 weeks. The mice were sacrificed after 2, 10 and 20 weeks at which time the capsules were removed and fixed in 4% paraformaldehyde for further analysis.

### Vaginal Cytology Collection and Analysis

To monitor animals for cycling, vaginal cytology was collected prior to ovariectomy and 7 days post-surgical recovery through animal endpoint. Daily, sterile saline was pipetted into the vaginal canal of mice using sterile glass pipettes and collected. Every other day longitudinal samples were imaged with a bright field microscope (Leica, Leica DMI3000B), where cells were identified, and samples were assigned a stage informed by the populations of cells imaged as defined by Cora et al ^24^. Estrus was defined by mouse that reached estrus stage for at least one day, whereas activity was classified by mice that were in metestrus or proestrus for at least one day. Stages were charted for individual mice to create a timeline for cycling.

### ELISAs for Hormone Analysis

Biweekly and terminal serum was collected from the mice via lateral tail vein or cardiac puncture respectively. Whole blood was stored at 4°C overnight to clot, and serum was isolated by spinning the samples at 2,000 g for 15 minutes. Serum samples were aliquoted and stored at -80°C until completion of the study. Samples were processed at the Ligand core at the University of Virginia. FSH quantification was performed using an ultrasensitive ELISA with a detectable range of 0.016-8 ng/mL (sensitivity of 0.016 ng/mL); The capture polyclonal antibody (guinea pig anti-mouse FSH; AFP-1760191; RRID: AB_2665512) and detection polyclonal antibody (rabbit anti-rat FSH-S11; AFP-C0972881; RRID: AB_2687903) were provided by the National Hormone and Peptide Program (NHPP). The HRP-conjugated polyclonal antibody (donkey anti-rabbit, AP182P; RRID: AB_92591) was purchased from EMD Millipore, Temecula, CA. The Mouse FSH reference preparation (mFSH-RP, AFP5308D; NHPP) is used as the assay standard. Estradiol quantification was performed using an estradiol ELISA kit (ALPCO, 11-ESTHU-E01) with a reportable range of 5-3200 pg/mL (sensitivity 5pg/ml) and progesterone quantification was performed using commercially available ELISA kit (Immuno-biological Laboratories Inc., IB79183) with a detectable limit of 0.1-40 ng/mL (sensitivity of 0.15 ng/ml).

### Embedding, sectioning and staining of histological samples

Samples were processed at the Histology Core at the University of Michigan Dental School. Implants were embedded in paraffin blocks and sectioned at a thickness of 5 µm, then every fourth slide was stained with hematoxylin (Abcam, ab220365) and eosin Y (Fisher Scientific, 2845). Histological slides were imaged at 10x and 20x using an upright brightfield microscope (Leica, DM1000LED).

### Stromal Cell Density Measurement and Follicle Counts

To quantify cellular density, 3 H+E sections were selected, an ROI (400 µm by 400 µm square region) was randomly selected, and the cell nuclei were counted using the cell counter function in Qupath. To control for tissue depth, slides were selected from the top 500 µm of the tissue, with the average depth per group ranging from 300 to 350 µm across conditions. Nuclei counts were normalized by area to calculate the cellular density per mm^2^.

To estimate the number of follicles per mm^3^ fresh pieces of ovarian tissue from each donor were embedded and sectioned and stained as described. Follicles in every eighth section were manually counted and annotated using QuPath software, with a total of 14-25 sections counted per donor. Densities were calculated by normalizing the follicle number to the tissue area with an estimated section thickness of 20 μm.

### Immunofluorescence Staining

Three sections at least 200 μm apart were selected for immunofluorescence staining. The slides were deparaffinized with xylenes (Thermo Fisher Scientific, X3P), rehydrated with a sequential ethanol wash and blocked with 0.03% hydrogen peroxide (Thermo Fisher Scientific, H324500). Next, the slides were treated using heated antigen retrieval in 0.1 M sodium citrate buffer (Thermo Fisher Scientific, J63950-AP) (90°C) for 20 minutes and blocked with normal goat serum 10% (ABCAM, NC0605700) and glycine solution (500M, BioRad, 1610717). Primary antibodies anti-pro and cleaved caspase 3 (rabbit anti-human, Novus, NB100-56113, RRID:AB_3073989) and anti-mixed lineage kinase domain-like protein (rat anti-human, Milipore, MABC1636, RRID: AB_2940818) were incubated overnight at 4°C. Samples then underwent a wash with tris buffered saline containing Tween-20 0.05% (Thermofisher, 85113) and Bovine Serum Albumin 0.1% (BSA, Thermofisher, MP219989910), followed by incubation with secondary anti-rat (Thermo Fisher Scientific, A48265TR, RRID: AB-2896334) and anti-rabbit (Thermo Fisher Scientific, A32731, RRID:AB_2633280) antibodies. The slides were mounted using prolong diamond mountant (Fisher Scientific, P36961) and imaged on a scanning laser confocal microscope (Leica,SP8) at 20x and 63x magnification.

### Immunofluorescence Image Processing

Using ImageJ, follicles were identified, and sections containing the oocyte and granulosa layers were selected. The mean grey value of the cas3 antibody signal in the identified follicle and immediate stroma (excluding the follicle) ROIs was collected. The mean grey value of the stroma was then normalized by the cellular density in the ROI to calculate the mean grey value per cell.

### Statistical Methods

Statistical analysis was performed using GraphPad Prism software. Data based on counts was analyzed using a chi-square analysis. Data was tested to confirm normality using Shapiro-Wilk test. Nonparametric or parametric one-way ANOVAs were used to compare in gel characteristics and cellular data, followed by Tukeys or Kruskal Wallace post hoc testing to distinguish differences across the gel conditions. Results were considered statistically different when p<0.05.

### The Use of Artificial Intelligence

To identify pores from other hydrogel impurities an AI image classifier (Weka) was trained using sample data images. Artificial intelligence was also used in the ideation of experiments or analysis of data, but was used to provide manuscript edits.

## Acknowledgements

The authors would like to acknowledge the University of Michigan Dental School Histology core and University of Virginia Ligand Core for their help with this work.

## Data Availability Statement

The data that support these findings of this study are available from the corresponding author upon request.

## Author Contributions

D.S., M.B. and A.S. conceived the project and designed the experiments. D.S., B.R, M.T, and S.T. fabricated microfluidic devices, and created gelatin microgels. D.S. carried material characterization, mechanical testing and diffusion experiments. D.S and A.S. performed animal surgeries. D.S., D.P., M.R., B.R., M.T., and S.T. conducted animal sample collection. D.S., B.R., M.T. and S.T. conducted staining experiments and analysis. D.S and A.S. drafted the manuscript with input from all authors. All authors discussed the results and approved the final manuscript.

## Funding

National Institutes of Health RO1- HD104173 (AS) National Institutes of Health R01-HD105018 (AS)

National Institutes of Health T32 Tissue Engineering and Regeneration Training Grant DE007057 (MAR, DIP, MAB)

National Institutes of Health T32 Training in Reproductive Biology Hd079342 (MAB, DSS)

Eunice Kennedy Shriver NICHD Z1A HD009005 (MAB)

---

**Supplemental Figure 1:**
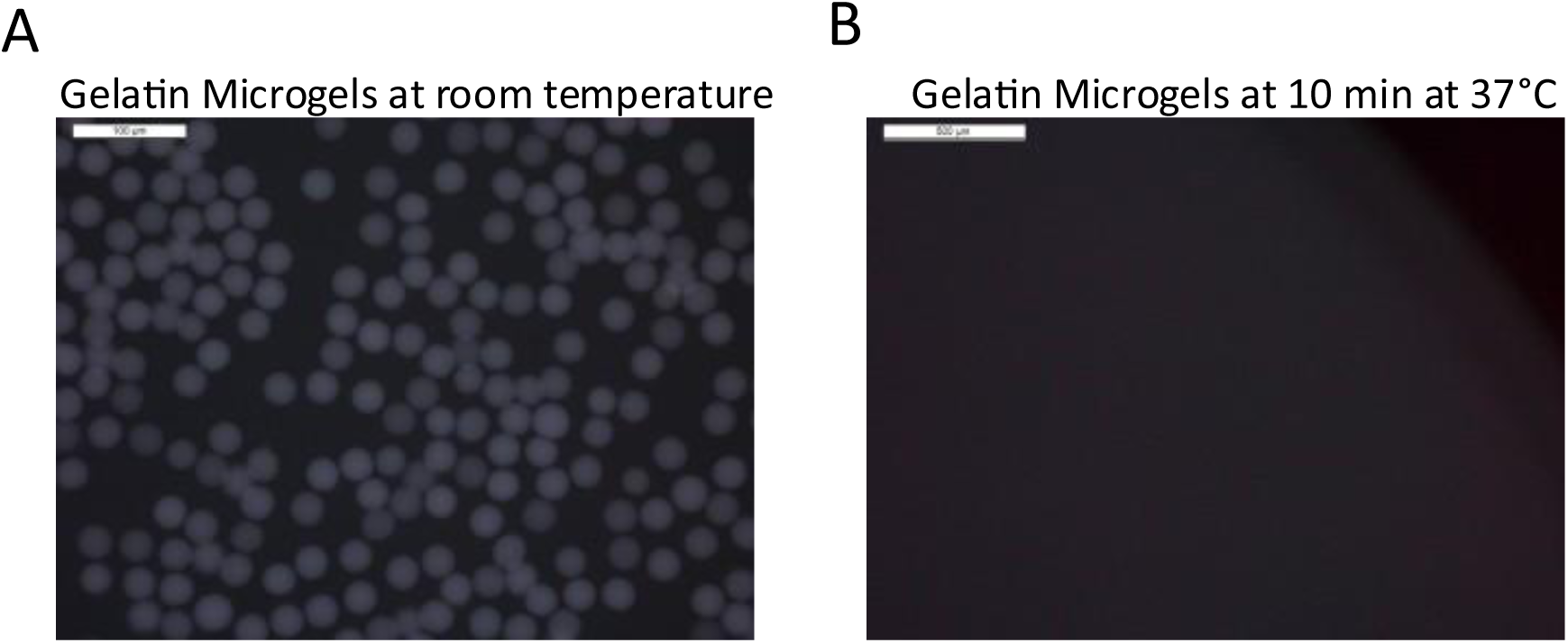
Microgel melting in solution. Gelati n microgels (A) at room temperature (20°C) and (B) 10 minutes at 37°C. Scale bar represents 100 μm (A) and 500 μm (B).

**Supplemental Figure 2:**
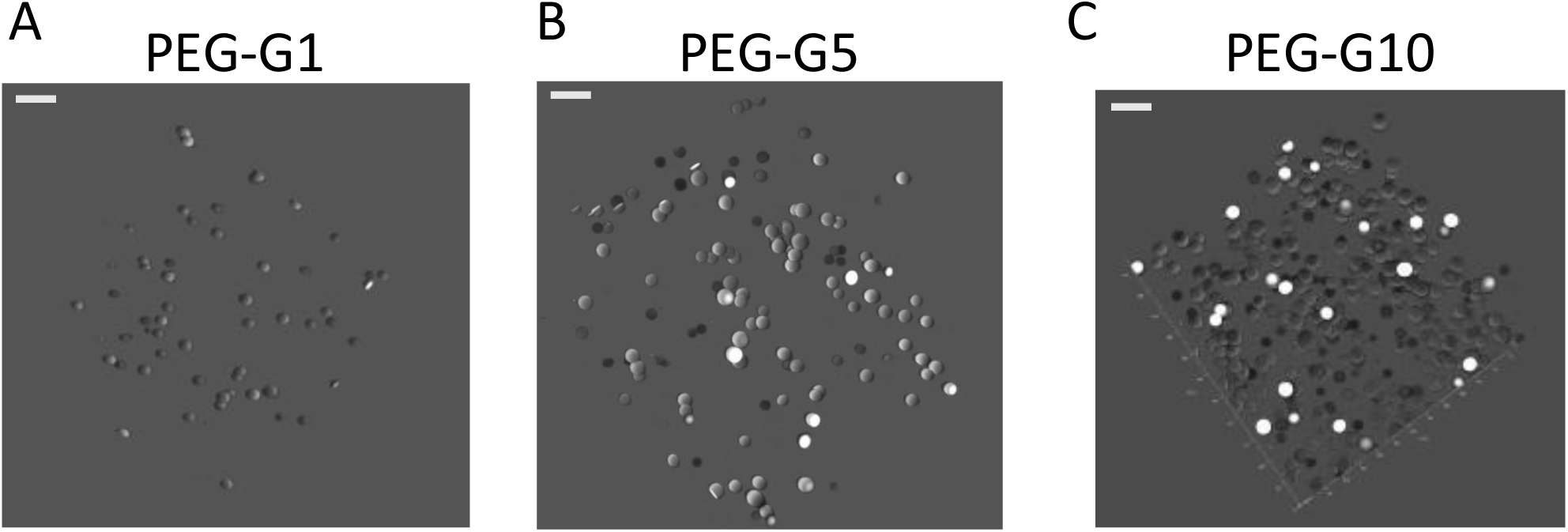
3D rendering of microgel distribution. 3D rendering of gelati n microgels distributed within a PEG hydrogel at (A) 1%, (B) 5% and (C) 10% volume/volume. Scale bar represents 100μm.

**Supplemental Figure 3:**
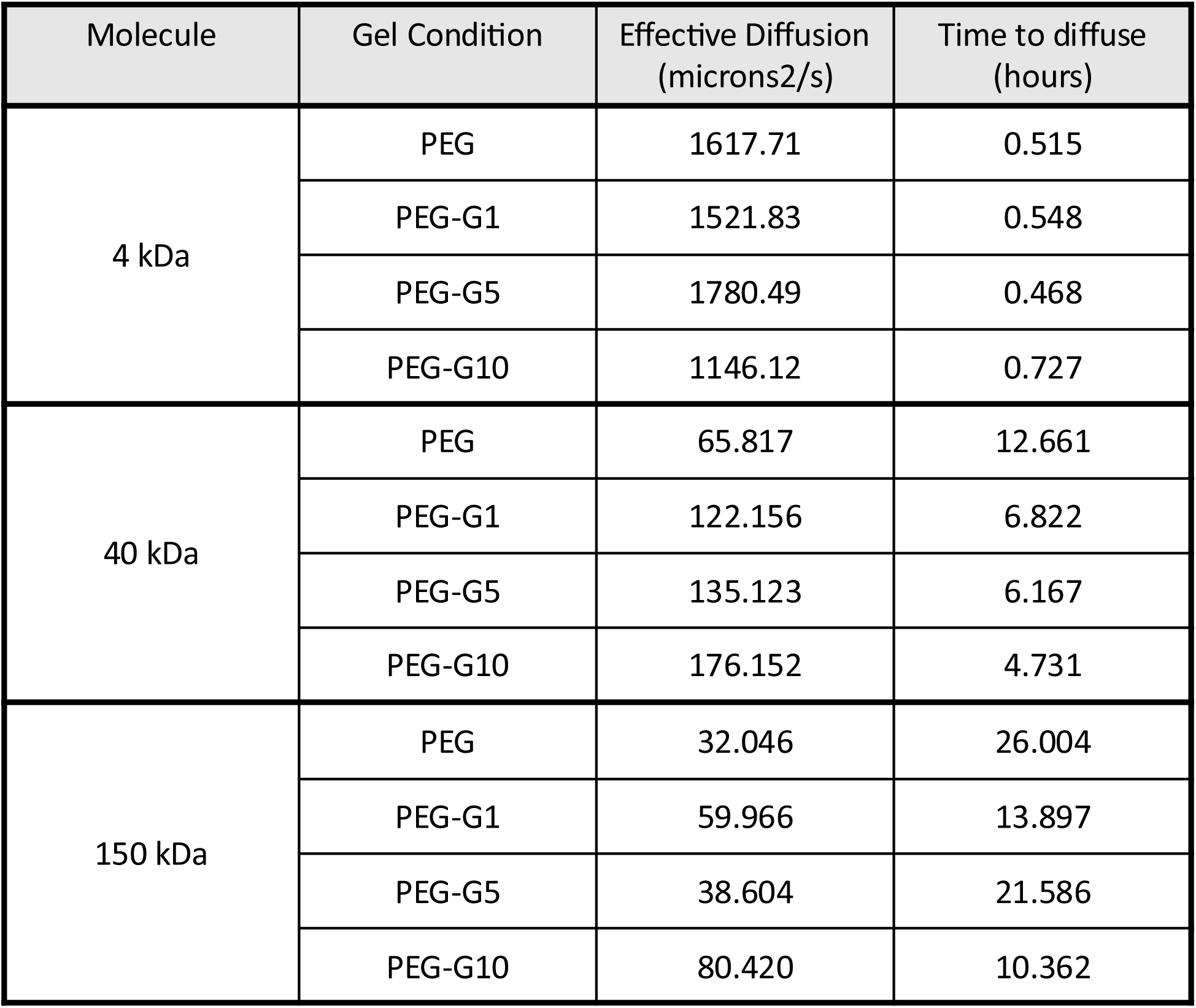
Calculated diffusivity and time to diffuse across the hydrogel. Calculated effective diffusion and time to diffuse through the hydrogel capsule.

**Supplemental Figure 4:**
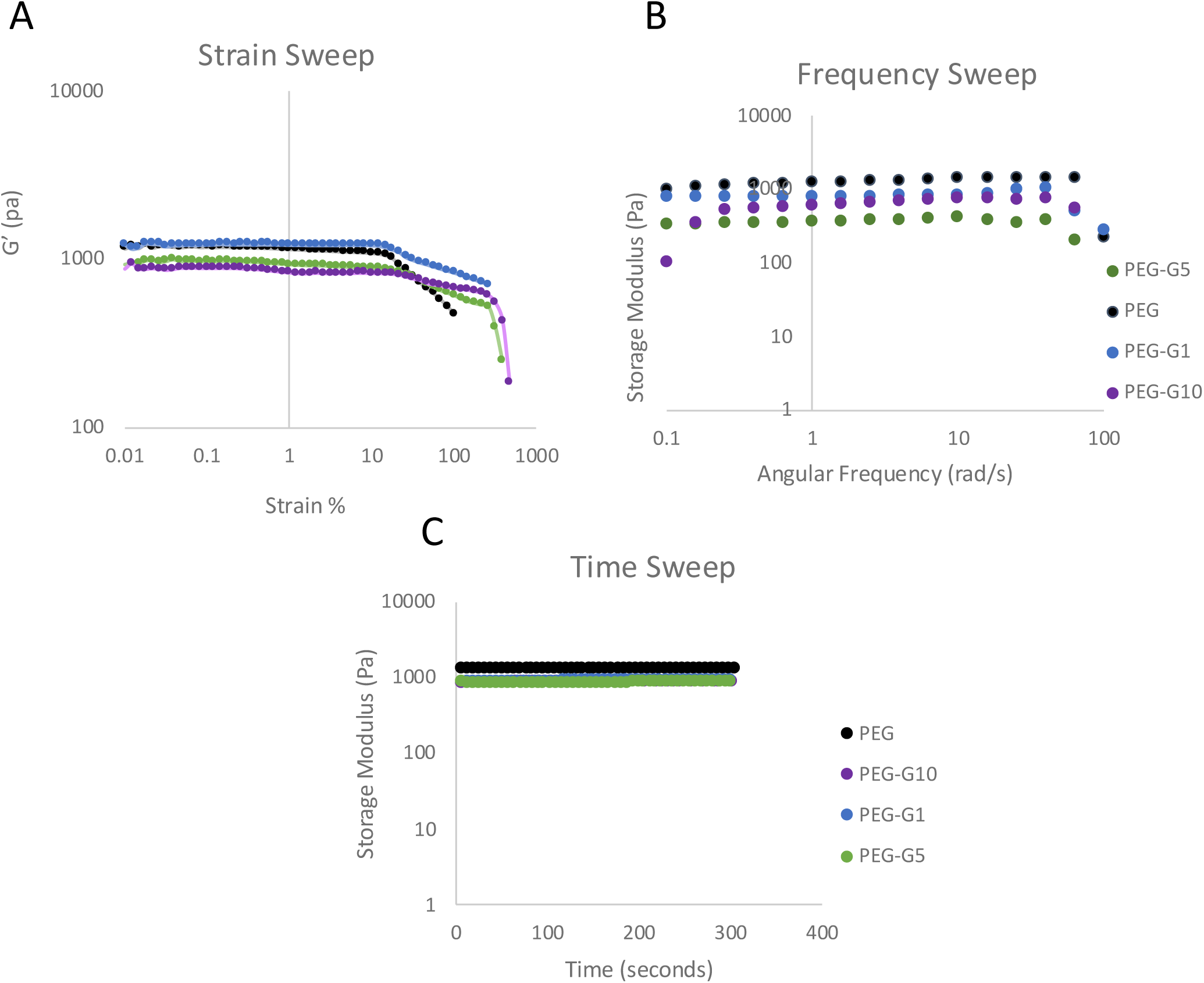
Example frequency and time sweeps. Representative (A) strain sweep, (B) frequency sweep and (C) time sweep at 37°C.

**Supplemental Figure 5:**
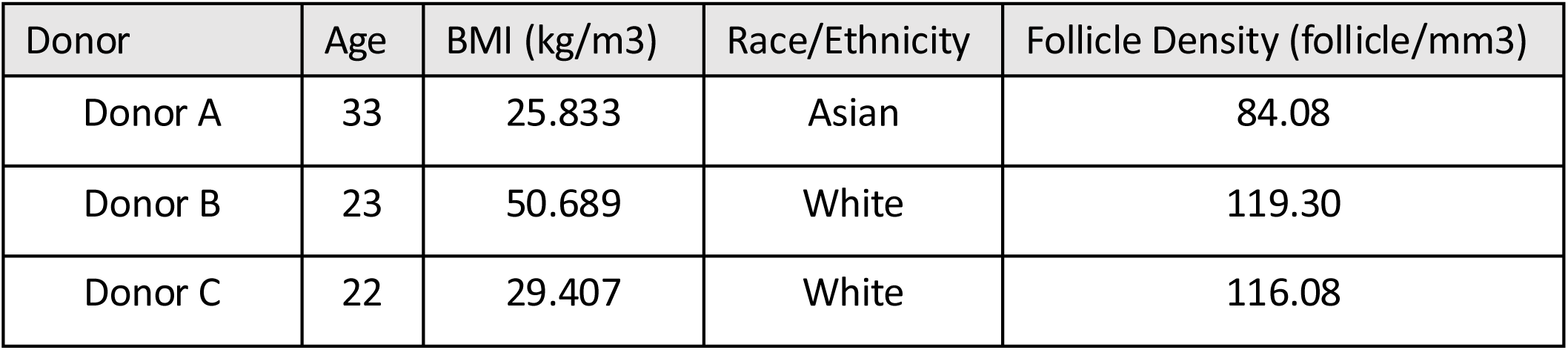
Donor information. Ovarian tissue donor data.

**Supplemental Figure 6:**
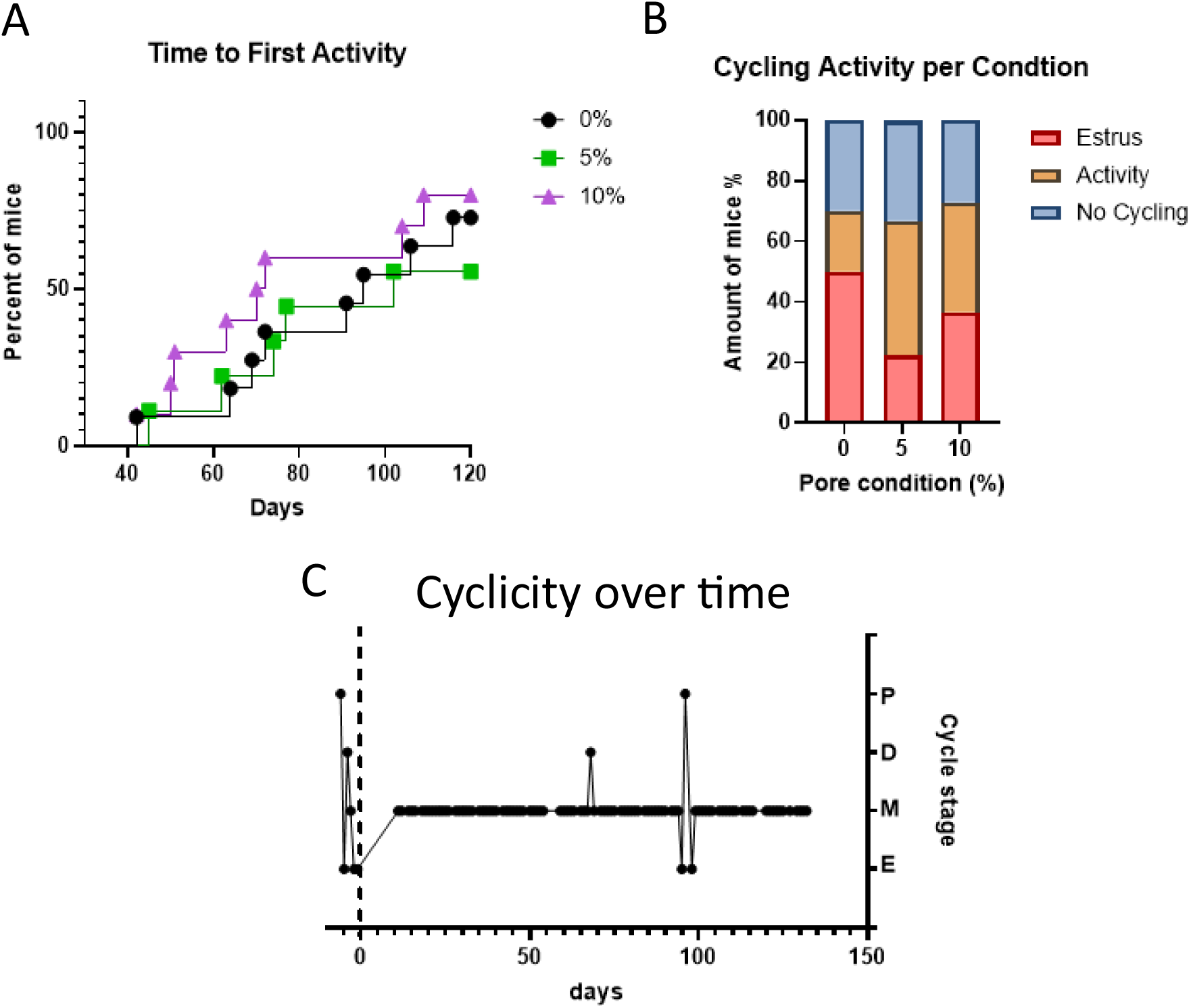
Estrus cyclicity. (A) Time to first cycling activity defined by proestrus or metestrus after prolonged diestrus. (B) Percent of mice that experienced estrus, only proestrus or metestrus (activity) and mice that did not deviate from diestrus. (C) Representative cytology plot.

**Supplemental Figure 7:**
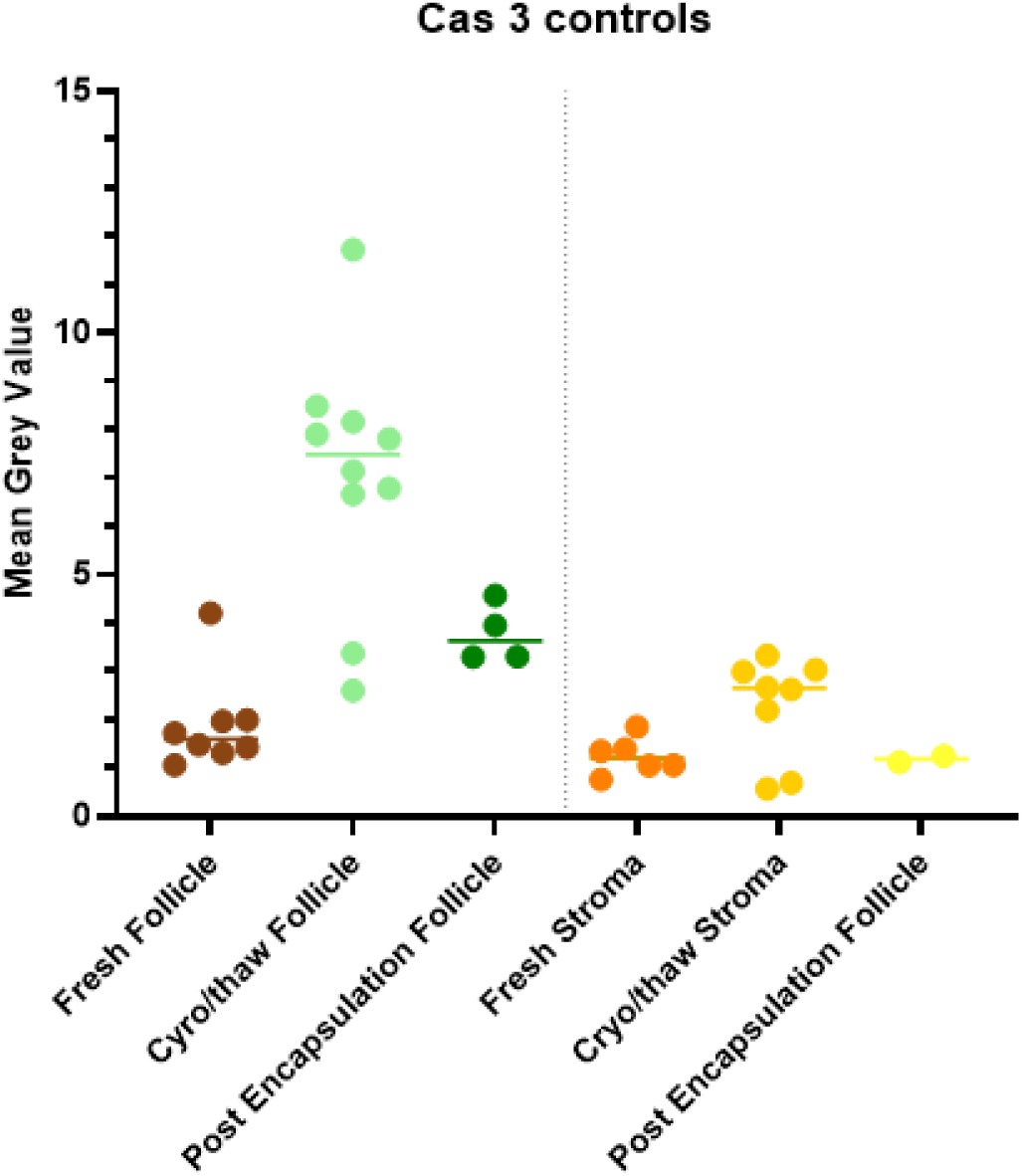
Cleaved caspase 3 staining controls. Comparison of Cleaved Caspase 3 staining in controls. Fresh tissue was fixed without freezing, Cryo/thaw tissue was fixed after cryopreservation and post encapsulation tissue was cryopreserved, encapsulated then fixed without implantation.

**Supplemental Figure 8:**
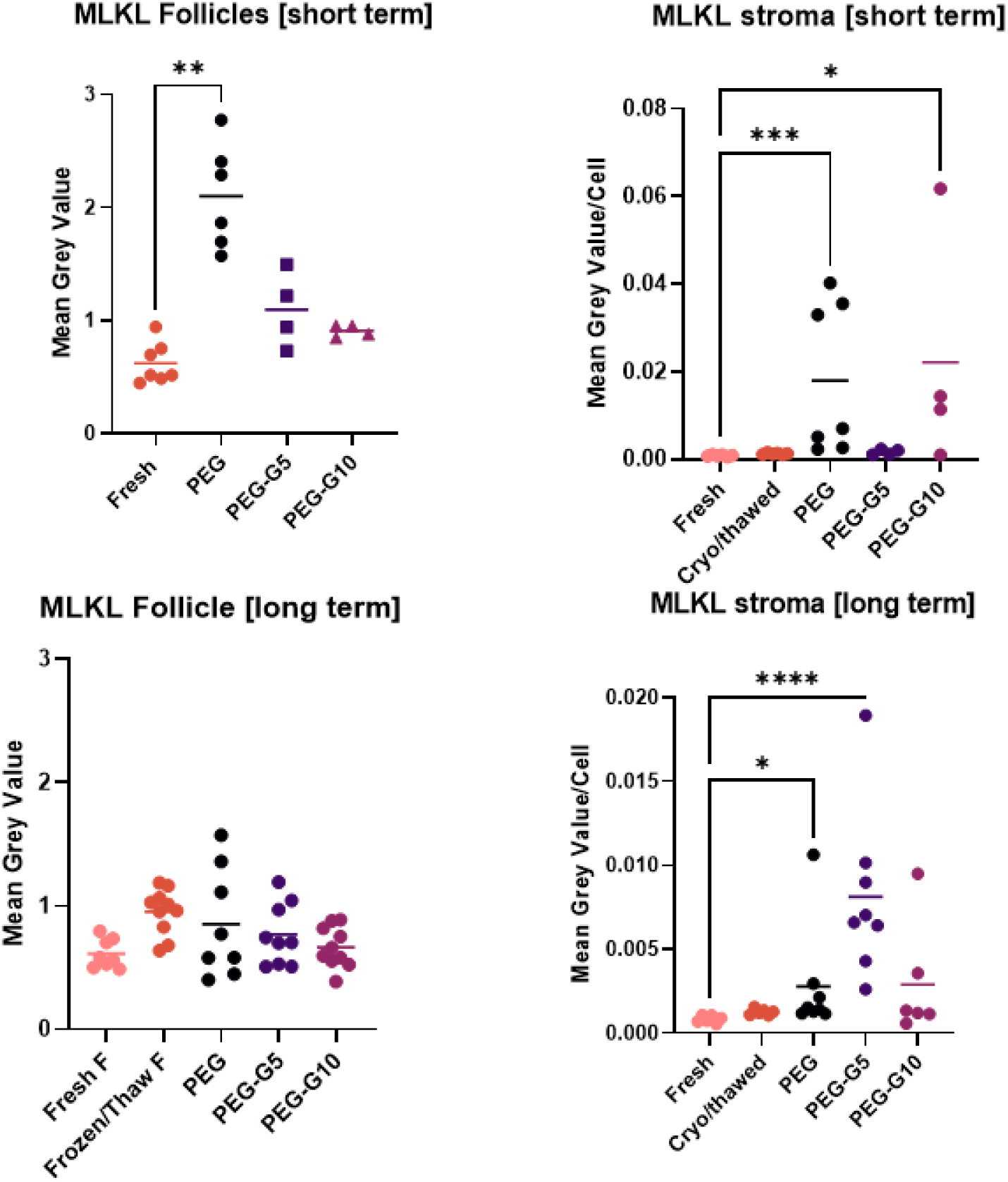
MLKL quantification. MLKL levels in short- and long-term encapsulated follicles and proximal stroma. One way ANOVA analysis with Tukey’s post hoc test.

## References

1. Siegel, R. L., Miller, K. D., Wagle, N. S. & Jemal, A. Cancer statistics, 2023. CA. Cancer J. Clin. 73, 17–48 (2023).

2. Chemaitilly, W. et al. Premature Ovarian Insufficiency in Childhood Cancer Survivors: A Report From the St. Jude Lifetime Cohort. J. Clin. Endocrinol. Metab. 102, 2242–2250 (2017).

3. Ruan, X. et al. Practice guideline on ovarian tissue cryopreservation and transplantation in the prevention and treatment of iatrogenic premature ovarian insufficiency. Maturitas 182, 107922 (2024).

4. Kolibianaki, E. E., Goulis, D. G. & Kolibianakis, E. M. Ovarian tissue cryopreservation and transplantation to delay menopause: facts and fiction. Maturitas 142, 64–67 (2020).

5. Dolmans, M.-M. et al. Transplantation of cryopreserved ovarian tissue in a series of 285 women: a review of five leading European centers. Fertil. Steril. 115, 1102–1115 (2021).

6. Kim, S. S. Assessment of long term endocrine function after transplantation of frozen-thawed human ovarian tissue to the heterotopic site: 10 year longitudinal follow-up study. J. Assist. Reprod. Genet. 29, 489–493 (2012).

7. Day, J. R. et al. Encapsulation of ovarian allograft precludes immune rejection and promotes restoration of endocrine function in immune-competent ovariectomized mice. Sci. Rep. 9, 16614 (2019).

8. Brunette, M. A. et al. Restoration of ovarian endocrine function with encapsulated immune-isolated human ovarian xenograft in ovariectomized mice. Sci. Adv. 12, eadx6348 (2026).

9. Brunette, M. A. et al. Human Ovarian Follicles Xenografted in Immunoisolating Capsules Survive Long Term Implantation in Mice. Front. Endocrinol. 13, (2022).

10. Ritger, P. L. & Peppas, N. A. A simple equation for description of solute release I. Fickian and non-fickian release from non-swellable devices in the form of slabs, spheres, cylinders or discs. J. Controlled Release 5, 23–36 (1987).

11. Meid, J. et al. Mechanical properties of temperature sensitive microgel/polyacrylamide composite hydrogels—from soft to hard fillers. Soft Matter 8, 4254–4263 (2012).

12. Fernandes, R. R., Andrade, D. E. V., Franco, A. T. & Negrão, C. O. R. The yielding and the linear-to-nonlinear viscoelastic transition of an elastoviscoplastic material. J. Rheol. 61, 893–903 (2017).

13. Day, J. R. et al. Immuno-Isolating Dual Poly(ethylene glycol) Capsule Prevents Cancer Cells from Spreading Following Mouse Ovarian Tissue Auto-Transplantation. Regen. Med. Front. 2019, e190006 (2019).

14. Slee, E. A., Adrain, C. & Martin, S. J. Executioner Caspase-3, -6, and -7 Perform Distinct, Non-redundant Roles during the Demolition Phase of Apoptosis*. J. Biol. Chem. 276, 7320–7326 (2001).

15. Chesnokov, M. S., Mamedova, A. R., Zhivotovsky, B. & Kopeina, G. S. A matter of new life and cell death: programmed cell death in the mammalian ovary. J. Biomed. Sci. 31, 31 (2024).

16. Markstrom, E., Svensson, Ec., Shao, R., Svanberg, B. & Billig, H. Survival factors regulating ovarian apoptosis -- dependence on follicle differentiation. https://doi.org/10.1530/rep.0.1230023 (2002) doi:10.1530/rep.0.1230023.

17. Kishi, H., Kitahara, Y., Imai, F., Nakao, K. & Suwa, H. Expression of the gonadotropin receptors during follicular development. Reprod. Med. Biol. 17, 11–19 (2018).

18. Candelaria, J. I., Rabaglino, M. B. & Denicol, A. C. Ovarian preantral follicles are responsive to FSH as early as the primary stage of development. J. Endocrinol. 247, 153–168 (2020).

19. Sokic, S., Christenson, M., Larson, J. & Papavasiliou, G. In Situ Generation of Cell-Laden Porous MMP-Sensitive PEGDA Hydrogels by Gelatin Leaching. Macromol. Biosci. 14, 731–739 (2014).

20. Kim, E. et al. Biomimetic composite gelatin methacryloyl hydrogels for improving survival and osteogenesis of human adipose-derived stem cells in 3D microenvironment. Mater. Today Bio 29, 101293 (2024).

21. Hwang, C. M. et al. Fabrication of three-dimensional porous cell-laden hydrogel for tissue engineering. Biofabrication 2, 035003 (2010).

22. Wang, W. Y. et al. Direct comparison of angiogenesis in natural and synthetic biomaterials reveals that matrix porosity regulates endothelial cell invasion speed and sprout diameter. Acta Biomater. 135, 260–273 (2021).

23. Day, J. R. et al. Immunoisolating poly(ethylene glycol) based capsules support ovarian tissue survival to restore endocrine function. J. Biomed. Mater. Res. A 106, 1381–1389 (2018).

24. Cora, M. C., Kooistra, L. & Travlos, G. Vaginal Cytology of the Laboratory Rat and Mouse: Review and Criteria for the Staging of the Estrous Cycle Using Stained Vaginal Smears. Toxicol. Pathol. 43, 776–793 (2015).

