## Supplemental Figures for "Microporous Immune-Isolating Capsule with Improved Diffusion for Restored Dynamic Bidirectional Hormone Signaling in a Murine Model of Premature Ovarian Insufficiency"

### Supplemental Figure 1:Microgel melting in solution

A

Gelatin Microgels at room temperature

B

Gelatin Microgels at 10 min at 37°C

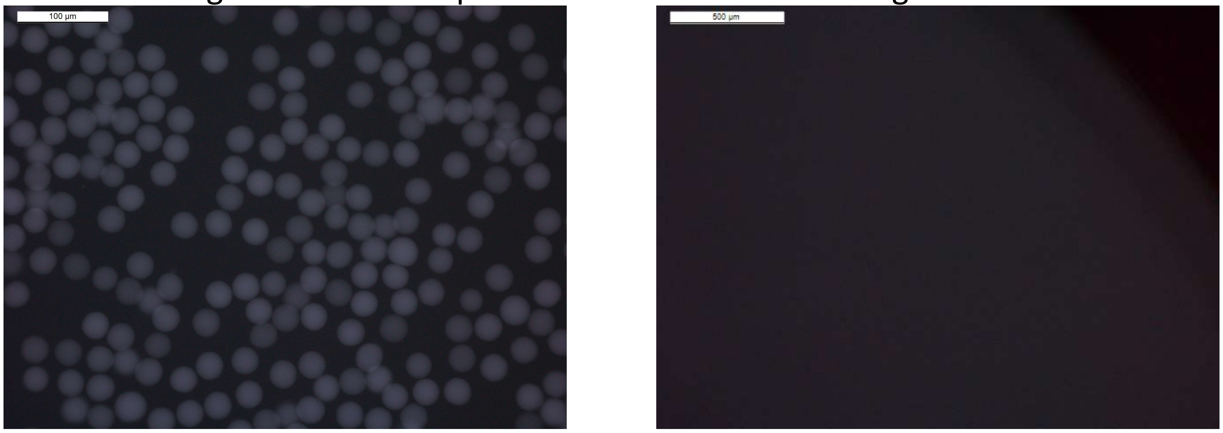


Supplemental Figure 1: Gelatin microgels (A) at room temperature (20°C) and (B) 10 minutes at 37°C. Scale bar represents 100 µm (A) and 500 µm (B).

### Supplemental Figure 2: 3D rendering of microgel distribution

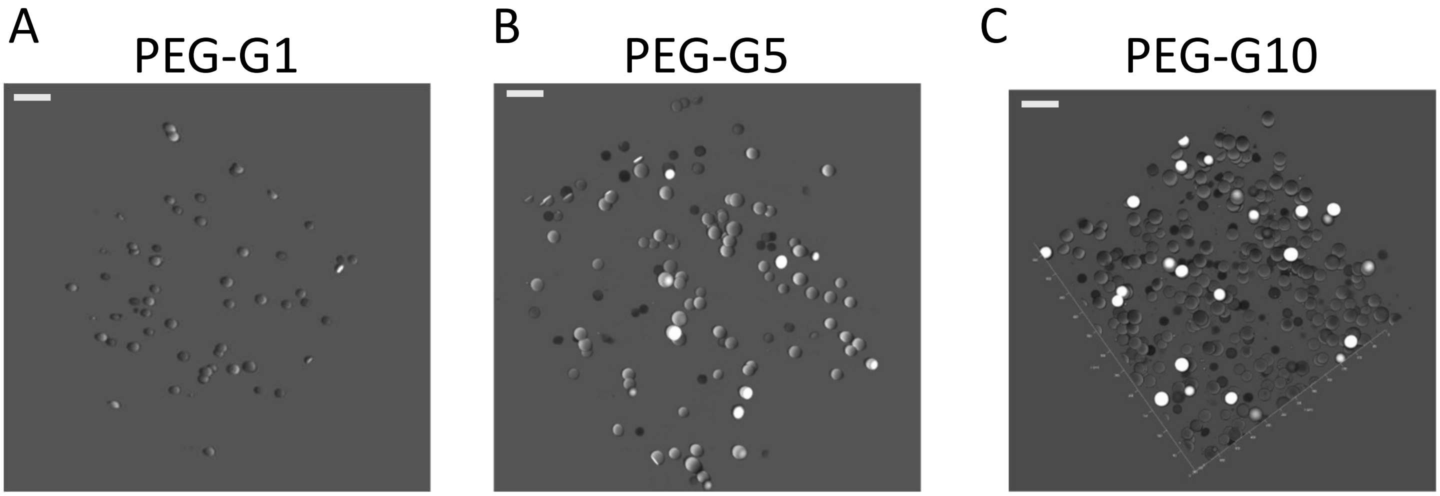


Supplemental figure 2: 3D rendering of gelatin microgels distributed within a PEG hydrogel at (A) 1%, (B) 5% and (C) 10% volume/volume. Scale bar represents 100µm.

### Supplemental Figure 3: Calculated diffusivity and time to diffuse across the hydrogel

| Molecule | Gel Condition | Effective Diffusion (microns2/s) | Time to diffuse (hours) |
|---|---|---|---|
| 4 kDa | PEG | 1617.71 | 0.515 |
| | PEG-G1 | 1521.83 | 0.548 |
| | PEG-G5 | 1780.49 | 0.468 |
| | PEG-G10 | 1146.12 | 0.727 |
| 40 kDa | PEG | 65.817 | 12.661 |
| | PEG-G1 | 122.156 | 6.822 |
| | PEG-G5 | 135.123 | 6.167 |
| | PEG-G10 | 176.152 | 4.731 |
| 150 kDa | PEG | 32.046 | 26.004 |
| | PEG-G1 | 59.966 | 13.897 |
| | PEG-G5 | 38.604 | 21.586 |
| | PEG-G10 | 80.420 | 10.362 |

Supplemental Figure 3: Calculated effective diffusion and time to diffuse through the hydrogel capsule.

### Supplemental Figure 4: Example frequency and time sweeps

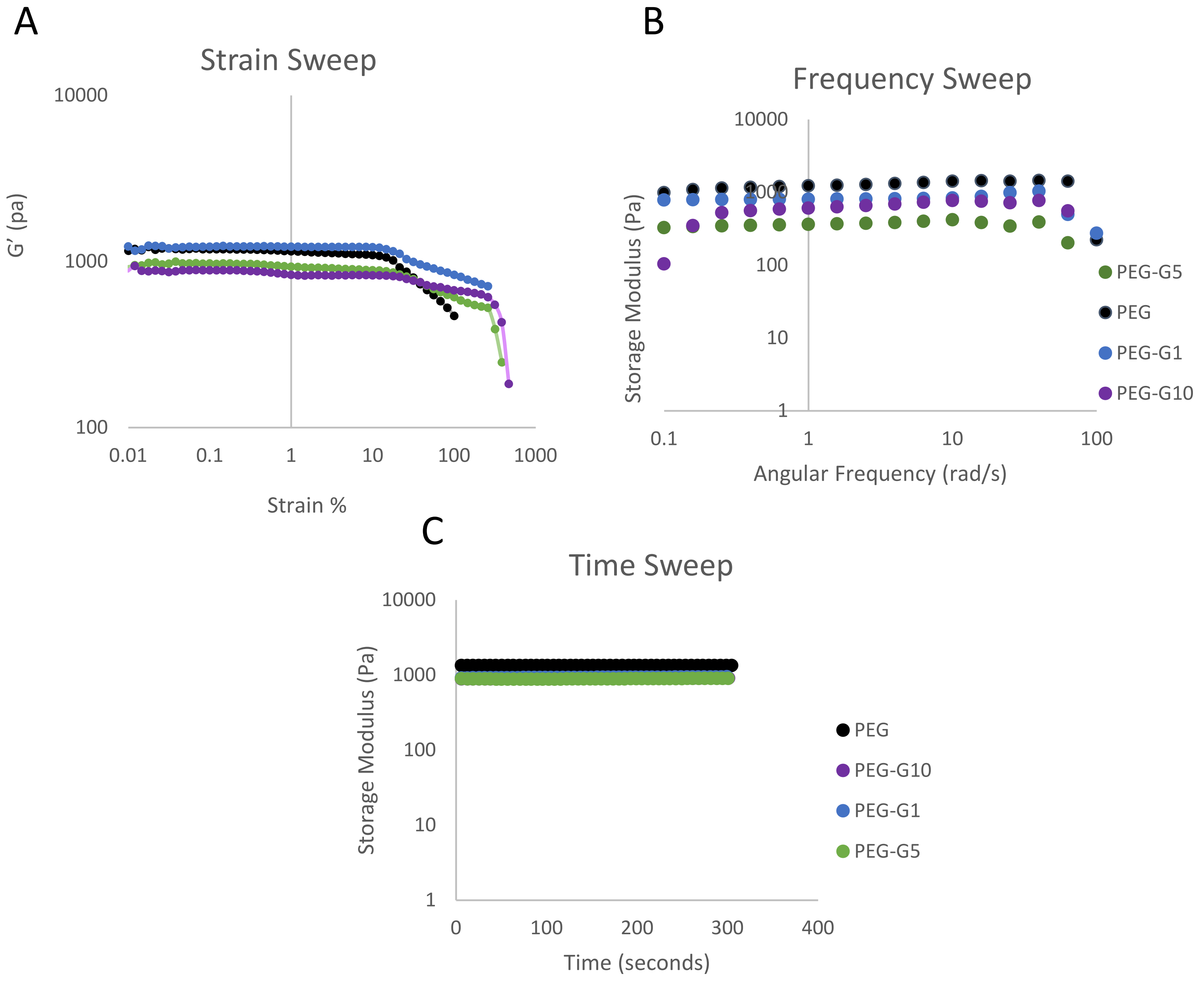


Supplemental Figure 4: Representative (A) strain sweep, (B) frequency sweep and (C) time sweep at 37°C.

### Supplemental Figure 5:Donor information

| Donor | Age | BMI (kg/m3) | Race/Ethnicity | Follicle Density (follicle/mm3) |
|---|---|---|---|---|
| Donor A | 33 | 25.833 | Asian | 84.08 |
| Donor B | 23 | 50.689 | White | 119.30 |
| Donor C | 22 | 29.407 | White | 116.08 |

Supplemental Figure 1: Ovarian tissue donor data.

### Supplemental Figure 6: Estrus cyclicity

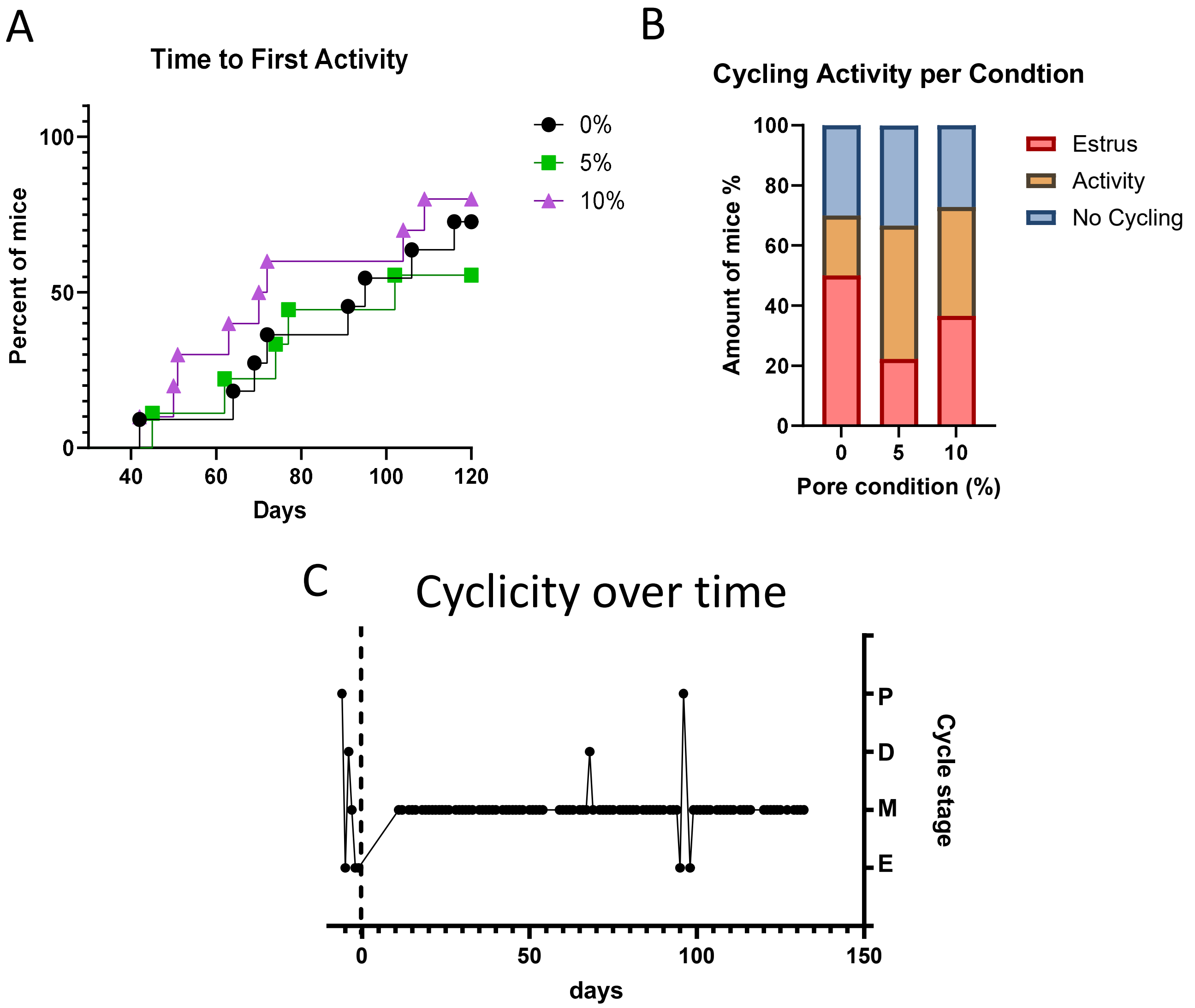


Supplemental Figure 6: (A) Time to first cycling activity defined by proestrus or metestrus after prolonged diestrus. (B) Percent of mice that experienced estrus, only proestrus or metestrus (activity) and mice that did not deviate from diestrus. (C) Representative cytology plot.

### Supplemental Figure 7: Cleaved caspase 3 staining controls

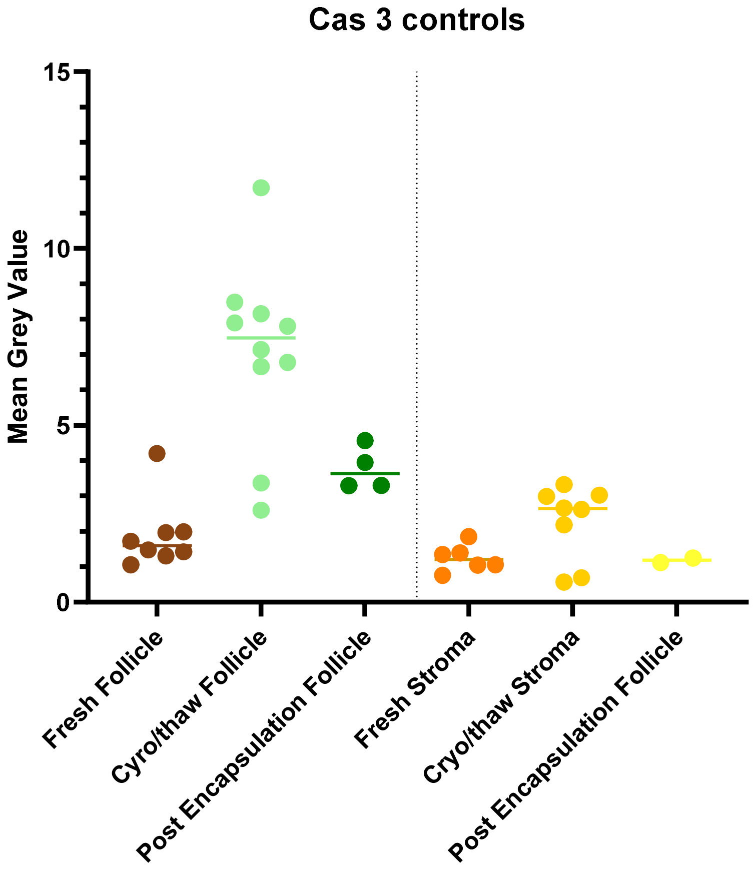


Supplemental Figure 7: Comparison of Cleaved Caspase 3 staining in controls. Fresh tissue was fixed without freezing, Cryo/thaw tissue was fixed after cryopreservation and post encapsulation tissue was cryopreserved, encapsulated then fixed without implantation.

### Supplemental Figure 8: MLKL quantification

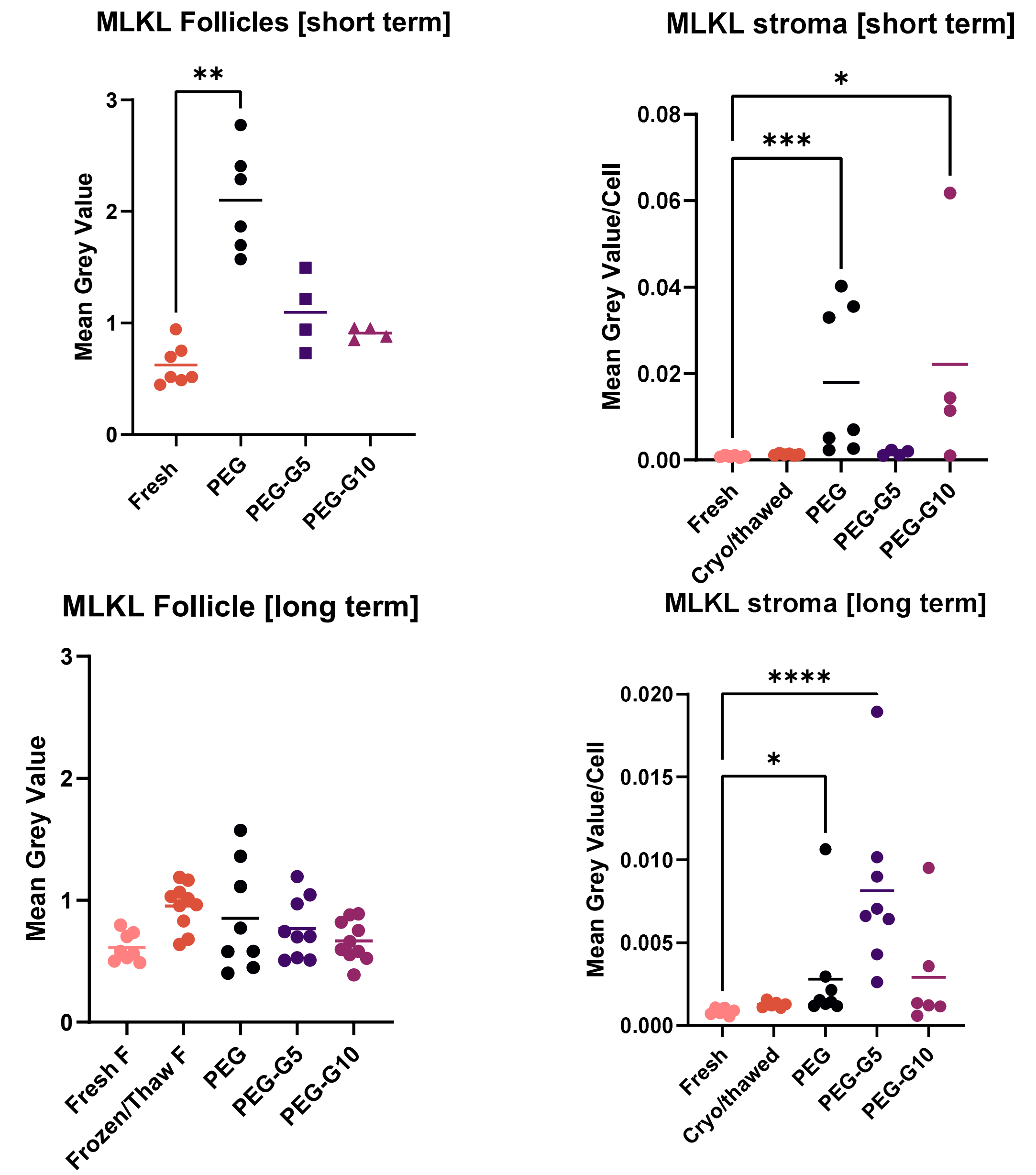


Supplemental Figure 8: MLKL levels in short- and long-term encapsulated follicles and proximal stroma. One way ANOVA analysis with Tukey's post hoc test.
